# Overcoming Attenuation Bias in Regressions using Polygenic Indices: A Comparison of Approaches

**DOI:** 10.1101/2021.04.09.439157

**Authors:** Hans van Kippersluis, Pietro Biroli, Rita Dias Pereira, Titus J. Galama, Stephanie von Hinke, S. Fleur W. Meddens, Dilnoza Muslimova, Eric A.W. Slob, Ronald de Vlaming, Cornelius A. Rietveld

## Abstract

Measurement error in polygenic indices (PGIs) attenuates the estimation of their effects in regression models. While this measurement error shrinks with growing Genome-wide Association Study (GWAS) sample sizes, the marginal returns to bigger sample sizes are rapidly decreasing. We analyze and compare two alternative approaches to reduce measurement error: Obviously Related Instrumental Variables (ORIV) and the PGI Repository Correction (PGI-RC). Through simulations, we show that both approaches outperform the typical (meta-analysis based) PGI in terms of bias and root mean squared error. Between families, the PGI-RC performs slightly better than ORIV, unless the prediction sample is very small (*N <* 1, 000), or when there is considerable assortative mating. Within families, ORIV is the default choice since the PGI-RC is not available in this setting. We verify the empirical validity of the simulations by predicting educational attainment (EA) and height in a sample of siblings from the UK Biobank. We show that applying ORIV between families increases the standardized effect of the PGI by 12% (height) and by 22% (EA) compared to a meta-analysis-based PGI, yet remains slightly below the PGI-RC estimates. Furthermore, within-family ORIV regression provides the tightest lower bound for the direct genetic effect, increasing the lower bound for the direct genetic effect on EA from 0.14 to 0.18, and for height from 0.54 to 0.61 compared to a meta-analysis-based PGI.

## Introduction

Genome-wide Association Studies (GWASs) have firmly established that, with few exceptions, most human (behavioral) traits are highly “polygenic” — that is, influenced by many individual genetic variants, each with a very small effect size (1; 2). A natural consequence of this has been the widespread adoption of so-called polygenic indices (PGIs), weighted sums aggregating the small effects of numerous genetic variants (single-nucleotide polymorphisms; SNPs), which enable out-of-sample genetic prediction of complex traits (3; 4; 5). It is common practice to meta-analyze GWAS summary statistics from as many samples as possible to foster the identification of genome-wide significant SNPs (1). Through the law of large numbers, this strategy has also proven to be very effective in reducing measurement error in the PGI and thus to boost the power and accuracy for analyses involving PGIs (4). PGIs are now able to explain a non-negligible proportion of the variance in health and behavioral traits (6).

Empirically constructed PGIs are nonetheless a noisy proxy for the true (latent) PGI because, amongst other reasons, the GWAS underlying the construction of the PGI is based on a finite sample (7; 8). As a result, the predictive power of today’s PGIs is still substantially smaller than the SNP-based heritability estimates, which constitute an upper bound for the predictive power of PGIs (9). For example, the current predictive power, or variance explained in the phenotype (*R*^2^), of the educational attainment (EA) PGI is about 12-16% (10), whereas the SNP-based heritability is estimated to be in the range 22-28% (e.g., 11; 12; 13). Importantly, the returns to GWAS sample size in terms of gaining predictive power of the PGI are rapidly decreasing (as described in e.g., 14; 15). For example, for a trait with a SNP-heritability of 25%, it takes a sample of ∼1 million to construct a PGI explaining 20%, but it would take a 7-fold increase in discovery sample size to achieve an *R*^2^ of 24%. Hence, reducing measurement error in the conventional way of meta-analyzing ever larger discovery samples has rapidly diminishing payoffs.

There is a burgeoning body of papers exploiting PGIs in medicine, biology and the social sciences. In many applications, the goal is to estimate the regression coefficient of the true latent PGI (i.e., the ‘additive SNP factor’), defined as the best linear predictor of the phenotype from the measured genetic variants (6). Taking the EA PGI as an example, a non-exhaustive list of applications includes (i) studies estimating the pathways through which the EA PGI influences lifetime outcomes including speech and reading skills (e.g., 16) or wealth accumulation (e.g., 17); (ii) studies that estimate direct genetic effects using within-family estimation to understand the mechanisms through which molecular differences translate into education differences (e.g., 18); (iii) studies that investigate gene-environment interplay in EA (e.g., 19; 20), and (iv) studies including a PGI as a control variable to reduce omitted variable bias or improve precision (e.g., 21). While there have been important advances in our understanding of how functional priors and optimizing construction methods can enhance the predictive power of a given PGI (e.g., 22; 23; 24), most existing applications still employ a (meta-analysis based) PGI without applying a correction for measurement error.

Recent studies have laid out the advantages of a measurement error correction and either suggested OLS estimates that have undergone some reasonable disattenuation (6; 25) or IV estimation (26) to deal with measurement error in the PGI. First, DiPrete et al. (26) suggested an instrumental variables (IV) approach to reduce measurement error in the PGI as a by-product of their Genetic Instrumental Variable (GIV) method. The intuition of the IV approach is simple. When we split the GWAS discovery sample into two, we can obtain two PGIs that both proxy the same underlying ‘true’ PGI. For example, when splitting the UK Biobank (UKB) at random into two discovery samples, both resulting PGIs approximate the same true latent PGI. Hence, theoretically, their correlation should be 1. However, in practice, their correlation will be smaller than 1 since the GWAS sample sizes used to construct these PGIs are finite and therefore each PGI will be subject to measurement error. In case the sources of measurement error are independent and the relative variance of measurement error in the PGI is the same across the two discovery samples, then the correlation between the two PGIs reveals the degree of measurement error. The IV approach in turn uses this information to correct (or ‘scale’) the observed association between the PGI and the outcome. Second, Becker et al. (6) developed an approach to disattenuate *between-family* estimates of the PGI on a trait, on the basis of external information of the trait’s SNP-based heritability (see also (25)). Intuitively, in this approach the SNP-based heritability is estimated in a first step, after which the coefficient of the PGI is re-scaled to match this SNP-based heritability in a second step. Since this approach was proposed alongside the introduction of the PGI repository project (6), we will refer to this approach as the *PGI repository correction*, or PGI-RC.

In this study, we use simulations and empirical analyses to compare the IV and PGI-RC approach to reduce attenuation bias in analyses involving PGIs. Our goal is to estimate a coefficient that is free from attenuation bias due to measurement error in a PGI. As such, we are not primarily interested in boosting the out-of-sample predictive power of a given PGI (in terms of e.g., the *R*^2^) for which ever-increasing GWAS discovery samples remain important. We are also not primarily interested in estimating the direct (or ‘causal’) effect of a PGI. PGIs typically capture not only effects of inherited variation (direct effects), but also effects of demography and relatives (so-called indirect genetic effects, 27). Therefore, when correcting for measurement error in a between-family design, the target parameter is the coefficient of the additive SNP-factor, which encompasses both direct as well as indirect genetic effects. In a within-family design, the target parameter is the direct genetic effect only.

In the simulations, we compare the performance of a meta-analysis based PGI (benchmark) to (1) results obtained using an Instrumental Variable (IV) approach (26), which relies on two PGIs constructed from two non-overlapping GWASs, and (2) the PGI repository correction (PGI-RC) (6). For the IV approach, we avoid the arbitrary choice of selecting one PGI as the independent variable and the other as IV by using the recently developed Obviously-Related Instrumental Variables (ORIV) technique (28). In our comparison of the benchmark, ORIV, and PGI-RC we consider various degrees of (i) genetic nurture (i.e., the effect of parental genotype on the child’s outcomes), (ii) assortative mating, and (iii) genetic correlation across discovery and prediction samples. Throughout the scenarios, we conduct between- as well as within-family analyses and pay specific attention to sample sizes for discovery and prediction samples. In turn, in the empirical application, we compare the benchmark, ORIV, and PGI-RC using data on height and educational attainment from the sibling sample of the UK Biobank (29).

We find that both the PGI-RC as well as ORIV outperform the benchmark in all scenarios. Comparing among them, the PGI-RC is particularly valuable in between-family analyses when the GWAS discovery sample is very small (i.e., when the predictive power of a PGI is low) or when there exists imperfect genetic correlation between discovery and prediction sample (i.e., when the effects of SNPs on the outcome are very different across the discovery and prediction sample). In the presence of assortative mating, however, the PGI-RC is both more biased and less precise than ORIV. Moreover, the PGI-RC tends to exhibit some bias in case of small (i.e., *N <* 1, 000) prediction samples, and its confidence intervals are wider than ORIV’s when incorporating the uncertainty induced by the fact that the PGI-RC requires an estimate of SNP-based heritability, which itself is subject to estimation error.

As such, the application of the PGI-RC or ORIV in empirical applications requires a careful assessment of the setting in which the correction is applied. From a practical point of view, it is a particular advantage of ORIV that it can be easily implemented in all standard statistical software packages (see https://github.com/geighei/ORIV for implementations in Stata and *R*). Moreover, ORIV does not require external information on the SNP-based heritability. This feature makes ORIV particularly attractive for within-family studies aiming to estimate direct genetic effects, because SNP-based heritability estimates do not solely capture direct genetic effects, but also incorporate indirect genetic effects from relatives, assortative mating and population stratification (e.g., 30). In other words, external information on the level of attenuation is typically not available when the interest is in direct genetic effects.^1^ Hence, ORIV is more flexible since it does not require external information on the ‘true’ level of disattenuation, which is hard to obtain in within-family settings or for traits that are subject to considerable assortative mating.

## Results

### Simulation study

To compare the performance of meta-analysis, ORIV, and the PGI-RC to estimate the standardized effect of the PGI on an outcome variable, we developed a general-purpose Python tool called GNAMES (Genetic-Nurture and Assortative-Mating-Effects Simulator). This tool allows users to efficiently simulate multi-generational genotype and phenotype data under genetic nurture (GN) effects and assortative mating (AM). The GNAMES tool itself, a description of its technical details, and a tutorial, are freely available on the following GitHub repository: https://github.com/devlaming/gnames.

In the simulations, we partition simulated genotype and phenotype data into three sets: two sets with equal sample size to perform non-overlapping GWASs (*discovery*) and one set to construct PGIs (*prediction*). For both sets of GWAS results, we construct a separate PGI. In addition, we also construct a PGI based on the meta analysis of the two GWASs, in line with the common practice of using the largest possible discovery sample to construct a PGI. Since the SNPs are simulated to be independent, the PGI simply equals the sum of the SNPs, weighted by the respective GWAS coefficients.

In our setup, every family has two children. Importantly, in the GWASs, data for only one sibling per family are used, and so the resulting PGIs are based on between-family GWAS results, as is common in the literature (but see (27) for a recent exception). The outcome of each simulation is a data file with the individual’s ID, the father and mother’s ID, a simulated outcome, two PGIs constructed from the two non-overlapping GWAS discovery samples, and the meta-analysis PGI. Further details of the simulation design are provided in Supplementary Information A.

The simulations are calibrated based on educational attainment (EA), but we show that our conclusions hold for traits with a different level of heritability in Supplementary Information C.1. The SNP-based heritability is approximately equal to 25% for EA in most samples (11; 12; 13). Hence, we fix the SNP-based heritability of the outcome at 25%. In turn, we use different settings that vary in terms of:

- The prediction sample – To create realistic variation in prediction sample sizes we vary *N*_*prediction*_ in the range (1,000; 2,000; 4,000; 8,000; 16,000). For example, a recent study on EA (32) employs prediction samples of similar sizes (the Dunedin Study (*N* = 810), the Environmental Risk Longitudinal Twin (E-Risk) Study (*N* = 1, 860), AddHealth (*N* = 5, 526), the Wisconsin Longitudinal Study (WLS, *N* = 7, 111) and the Health and Retirement Study (HRS, *N* = 8, 546)).
- The size of the GWAS discovery sample – As described in (7; 8), the predictive power of a PGI mainly depends on the variance explained by each SNP, and the ratio of the GWAS discovery sample to the number of SNPs. In particular, under the assumption that all SNPs explain an equal proportion of the SNP-based heritability, De Vlaming et al. (33, equation 2) approximate the predicted *R*^2^ of a given PGI as

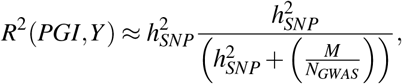

where 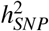 is the SNP-based heritability, *M* is the number of SNPs and *N*_*GWAS*_ denotes the GWAS discovery sample size. To maintain a manageable simulation space, we hold the number of SNPs fixed at *M* = 5, 000, and set 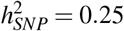. In order to simulate realistic levels of predictive power for a PGI, we use *N*_*GWAS*_ in the range (2,000; 4,000; 8,000; 16,000; 32,000). These values for the GWAS discovery samples then respectively generate an expected *R*^2^ of 2.3% (i.e., close to the PGI performance in the first EA GWAS, EA1, (34)), 4.2% (∼EA2, (12)), 7.1%, 11.1% (∼EA3, (35)) and 15.4% (∼EA4, (10)).^2^
- The presence of genetic nurture – We hold 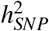 (i.e., the SNP-heritability) constant at 0.25 in each generation. In the presence of genetic nurture, the estimated SNP-based heritability is however a combination of direct genetic effects and genetic nurture (e.g., 31): 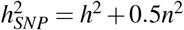, where *h*^2^ denotes the phenotypic variance accounted for by direct genetic effects, and *n*^2^ denotes the phenotypic variance accounted for by genetic nurture. In scenarios without genetic nurture, we fix the SNP-based heritability at 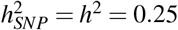. In scenarios with genetic nurture, we set *h*^2^ = 0.2 and *n*^2^ = 0.1 such that the genetic nurture is half the size of the direct genetic effect. This parametrization is again loosely following empirical evidence for EA, where the indirect genetic effect represents roughly half of the additive SNP factor (e.g., 10; 18).
- The degree of assortative mating – We vary assortative mating on the outcome variable (i.e., the phenotypic correlation between mates) between 0 and 1 in increments of 0.25.
- The genetic correlation – We vary the genetic correlation between the GWAS discovery samples and the prediction sample between 0.25 and 1 in increments of 0.25, where the two GWAS discovery samples have perfect genetic correlation. Additionally, we vary the genetic correlation between discovery sample 1 and discovery sample 2 from 0.25 to 1 in increments of 0.25, where we maintain a perfect genetic correlation between discovery sample 2 and the prediction sample.

For each scenario we perform 100 simulation replications (“runs”), and we average the results over these runs. In all scenarios, we estimate (i) a linear regression of the outcome on the meta-analysis PGI; (ii) ORIV, on basis of a two-stage least squares (2SLS) regression where we simultaneously use both independent PGIs as instrumental variables for each other; and (iii) the PGI-RC where we scale the coefficient obtained under (i) by the estimated SNP-based heritability. The SNP-based heritability is estimated in the prediction sample using MGREML (Multivariate Genome-based Restricted Maximum Likelihood, 36). In these GREML analyses, individuals with a genetic relatedness larger than 0.05 are excluded. The default PGI-RC estimator ignores the estimation error in the estimation of the scaling factor (as acknowledged on p. 14 of the Supplementary Information of 6). This may lead to anti-conservative standard errors, and in the univariate case the true standard error is equal to the standard error of the square root of the estimated SNP-heritability. We will therefore present both the default PGI-RC (“PGI-RC (Default)”) application, as well as the PGI-RC that incorporates the uncertainty in the scaling factor (“PGI-RC (GREML unc.)”). Details of the estimation procedures can be found in Methods.

Our main evaluation criterion is the resulting point estimate and 95% confidence interval of the estimated coefficient, to compare the bias of a particular method from the known true coefficient. However, in order to balance bias as well as precision, in Supplementary Information B we additionally present the root mean squared error, which is a function of both the bias as well as the variance.

#### Variation in prediction sample size

Figure 1 shows the coefficient estimates and their 95% confidence intervals for a meta-analysis PGI, ORIV and the PGI-RC for varying sample sizes of the prediction sample. The estimates derive from our baseline scenario (no genetic nurture, no assortative mating) and where the GWAS sample size is held constant such that the resulting meta-analysis PGI roughly corresponds to EA4 (i.e., *R*-squared ∼ 15.4%). Since we simulated a scenario without genetic nurture and assortative mating, the between-family (panel 1a) and within-family (panel 1b) analyses target the same coefficient (i.e., a SNP-based heritability of 0.25, or a standardized coefficient of 0.5; see Methods).

**Figure 1.**
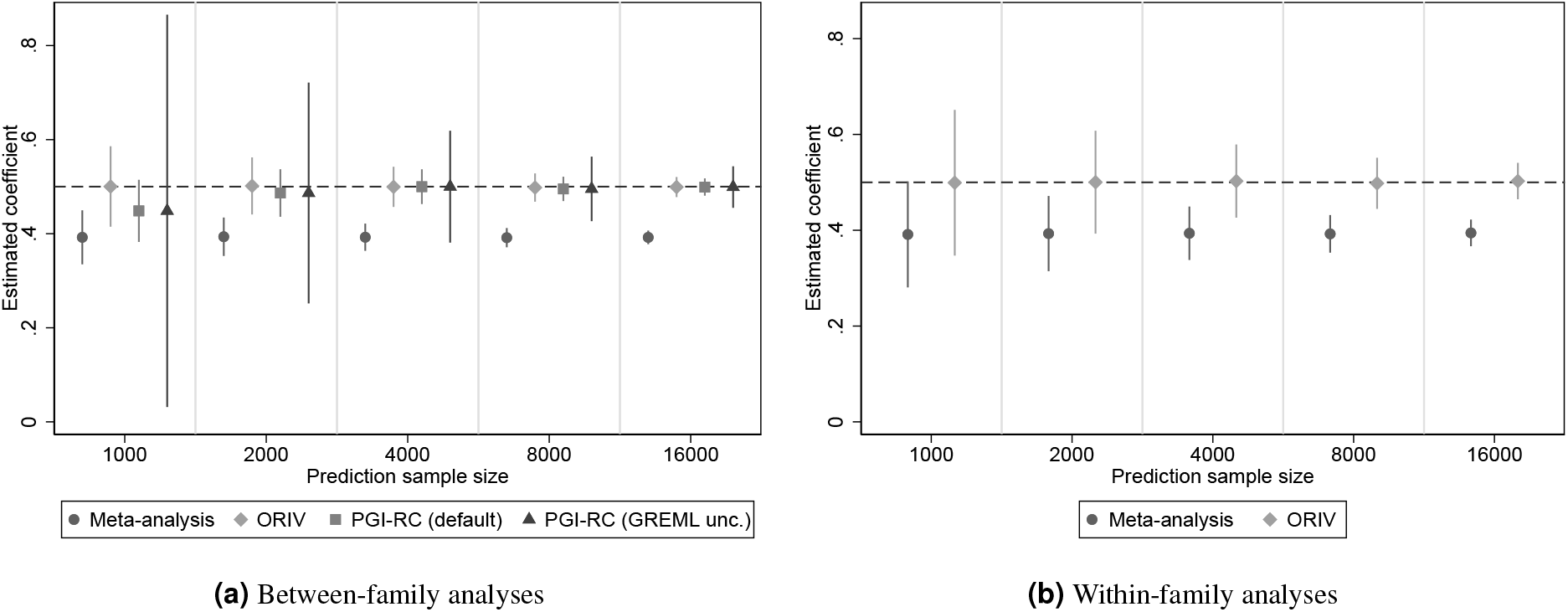
Estimated coefficients for the Polygenic Index (PGI) and their corresponding 95% confidence intervals using meta-analysis (circles), Obviously Related Instrumental Variables (ORIV, rhombuses), the default PGI-RC procedure (squares), and the PGI-RC procedure taking into account uncertainty in the GREML estimates (triangles) in the baseline scenario (no genetic nurture, no assortative mating) and holding constant the GWAS discovery sample such that the resulting meta-analysis PGI has an *R*^2^ of 15.4%. The horizontal dashed line represents the true coefficient. The simulation results are based on 100 replications.

Judging from the point estimates, in a between-family setting, ORIV and the PGI-RC clearly outperform a meta-analysis PGI, with limited differences between them in this scenario. An increasing prediction sample size shrinks the confidence intervals but leaves the coefficients largely unaffected. One notable exception is that for a small prediction sample size (*N* ≤ 1, 000), the PGI-RC slightly underestimates the true coefficient, because the estimated GREML SNP-heritability that is used in the PGI-RC is measured with noise in such a small sample. It is also in this setting with a relatively small prediction sample that the uncertainty in the PGI-RC (GREML unc.) is considerably larger than suggested in the default application PGI-RC (default). With a relatively large GWAS sample, the RMSE for ORIV tends to be slightly smaller than for the PGI-RC (see Table 4 in Supplementary Information B).

Within-families, the PGI-RC is not available. Confidence intervals are clearly larger within-families than between-families since the variation in the PGI is more limited, but again shrink with an increasing prediction sample size. Similar to the between-family analysis, within-families ORIV is consistent and clearly outperforms a meta-analysis based PGI in this scenario.

#### Variation in GWAS sample size

Figure 2 shows the corresponding figure from the same scenario but now holding the prediction sample size constant at *N* = 16, 000 and varying the GWAS sample size such that the *R*-squared of the meta-analysis PGI varies from 2.3% (EA1) to 15.4% (EA4). Whereas a meta-analysis PGI clearly benefits from an increased GWAS sample size, for a relatively large prediction sample size of *N* = 16, 000 both ORIV and the PGI-RC are approximately unbiased irrespective of the GWAS sample size. The PGI-RC shows a narrower confidence interval compared with ORIV, even when taking into account uncertainty in the GREML estimates, with the difference becoming smaller for larger GWAS discovery samples (see also Table 5 in Supplementary Information B).

**Figure 2.**
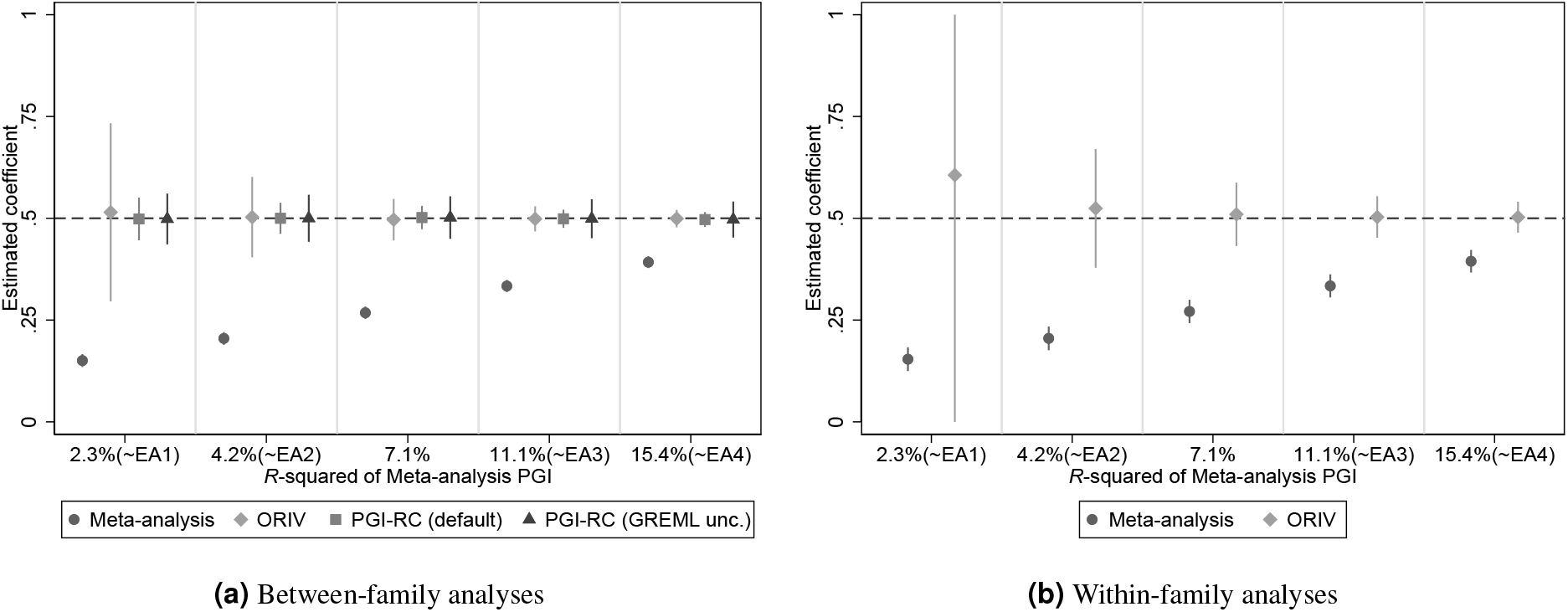
Estimated coefficients for the Polygenic Index (PGI) and their corresponding 95% confidence intervals using meta-analysis (circles), Obviously Related Instrumental Variables (ORIV, rhombuses), the default PGI-RC procedure (squares), and the PGI-RC procedure taking into account uncertainty in the GREML estimates (triangles) in the baseline scenario (no genetic nurture, no assortative mating) and holding constant the prediction sample at *N* = 16, 000. The confidence interval for EA1 in the within-family analysis extends beyond the displayed range (−0.08 – 1.29). The dashed line represents the true coefficient. The simulation results are based on 100 replications.

#### Variation in both prediction and GWAS sample size

In Table 1 we vary both the GWAS as well as the prediction sample size simultaneously. In particular, we compare the performance of a relatively small GWAS sample size, calibrated to be resulting in an *R*-squared of 4.2% (roughly EA2) for different prediction sample sizes, and a relatively large GWAS sample size, calibrated to be resulting in an *R*-squared of 15.4% (roughly EA4), again for different prediction sample sizes. As expected, when the GWAS sample size increases, the coefficient of a meta-analysis PGI comes closer to the true value (0.5), but even for a relatively large GWAS sample size it is still considerably biased. As before, an increasing prediction sample size does not decrease the bias in the coefficient of a meta-analysis PGI, but only shrinks the confidence interval.

**Table 1.**
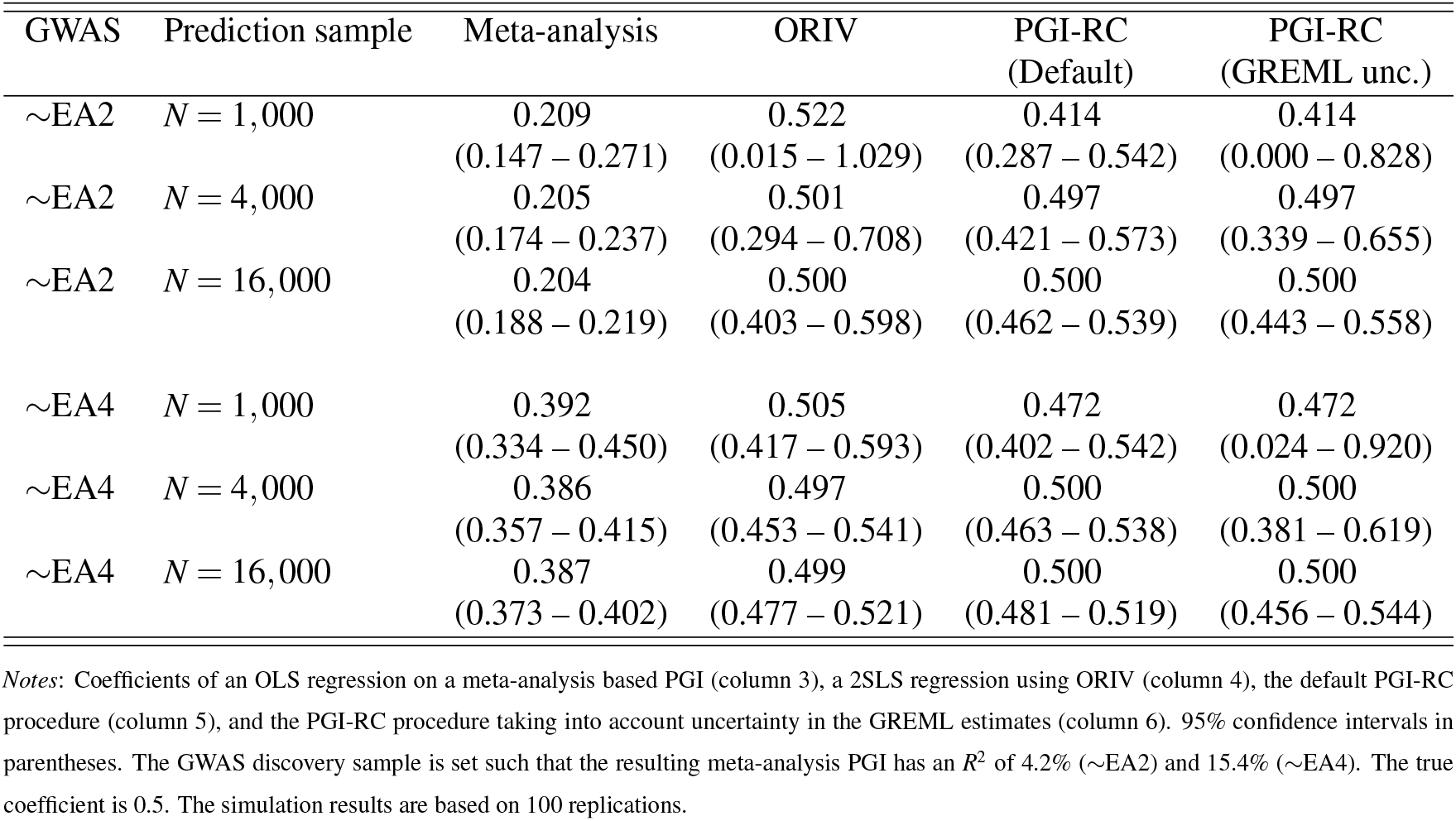
Estimated coefficients for the Polygenic Index (PGI) and their corresponding 95% confidence intervals using meta-analysis, Obviously Related Instrumental Variables (ORIV), and the PGI-RC procedure in the baseline scenario (no genetic nurture, no assortative mating; between-family analyses only).

| GWAS | Prediction sample | Meta-analysis | ORIV | PGI-RC<br>(Default) | PGI-RC<br>(GREML unc.) |
| --- | --- | --- | --- | --- | --- |
| ~EA2 | $N = 1,000$ | 0.209<br>(0.147 – 0.271) | 0.522<br>(0.015 – 1.029) | 0.414<br>(0.287 – 0.542) | 0.414<br>(0.000 – 0.828) |
| ~EA2 | $N = 4,000$ | 0.205<br>(0.174 – 0.237) | 0.501<br>(0.294 – 0.708) | 0.497<br>(0.421 – 0.573) | 0.497<br>(0.339 – 0.655) |
| ~EA2 | $N = 16,000$ | 0.204<br>(0.188 – 0.219) | 0.500<br>(0.403 – 0.598) | 0.500<br>(0.462 – 0.539) | 0.500<br>(0.443 – 0.558) |
| ~EA4 | $N = 1,000$ | 0.392<br>(0.334 – 0.450) | 0.505<br>(0.417 – 0.593) | 0.472<br>(0.402 – 0.542) | 0.472<br>(0.024 – 0.920) |
| ~EA4 | $N = 4,000$ | 0.386<br>(0.357 – 0.415) | 0.497<br>(0.453 – 0.541) | 0.500<br>(0.463 – 0.538) | 0.500<br>(0.381 – 0.619) |
| ~EA4 | $N = 16,000$ | 0.387<br>(0.373 – 0.402) | 0.499<br>(0.477 – 0.521) | 0.500<br>(0.481 – 0.519) | 0.500<br>(0.456 – 0.544) |

*Notes:* Coefficients of an OLS regression on a meta-analysis based PGI (column 3), a 2SLS regression using ORIV (column 4), the default PGI-RC procedure (column 5), and the PGI-RC procedure taking into account uncertainty in the GREML estimates (column 6). 95% confidence intervals in parentheses. The GWAS discovery sample is set such that the resulting meta-analysis PGI has an $R^2$ of 4.2% (~EA2) and 15.4% (~EA4). The true coefficient is 0.5. The simulation results are based on 100 replications.

**Table 2.** Results of the OLS and IV regressions explaining (residualized and standardized) educational attainment.

|  | OLS<br>(UKB) | OLS<br>(23andMe) | OLS<br>(Meta-analysis) | ORIV<br>(2-sample) | ORIV<br>(Split-sample) | PGI-RC<br>(default) | PGI-RC<br>(GREML unc.) |
| --- | --- | --- | --- | --- | --- | --- | --- |
| **Between-family results** |  |  |  |  |  |  |  |
| Polygenic index | 0.258***<br>(0.005) | 0.218***<br>(0.005) | 0.276***<br>(0.005) | 0.337***<br>(0.007) | 0.323***<br>(0.007) | 0.394***<br>(0.007) | 0.394***<br>(0.024) |
| First-stage estimate |  |  |  | 0.498***<br>(0.005) | 0.489***<br>(0.005) |  |  |
| First-stage $F$-statistic | | | | 11,918.98 | 11,061.02 | | |
| Incremental $R^2$ | 6.7% | 4.7% | 7.6% | | | | |
| Family fixed effects | NO | NO | NO | NO | NO | NO | NO |
| $N$ | 35,282 | 35,282 | 35,282 | 35,282 | 35,282 | 35,282 | 35,282 |
| **Within-family results** |  |  |  |  |  |  |  |
| Polygenic index | 0.124***<br>(0.009) | 0.115***<br>(0.009) | 0.142***<br>(0.009) | 0.184***<br>(0.012) | 0.170***<br>(0.013) |  |  |
| First-stage estimate |  |  |  | 0.460***<br>(0.006) | 0.436***<br>(0.006) |  |  |
| First-stage $F$-statistic | | | | 6,068.34 | 5,184.87 | | |
| Incremental $R^2$ | 1.5% | 1.3% | 2.0% | | | | |
| Family fixed effects | YES | YES | YES | YES | YES |  |  |
| $N$ | 35,282 | 35,282 | 35,282 | 35,282 | 35,282 | | |

**Table 3.** Results of the OLS and IV regressions explaining (residualized and standardized) height.

|  | OLS<br>(UKB) | OLS<br>(GIANT) | OLS<br>(Meta-analysis) | ORIV<br>(2-sample) | ORIV<br>(Split-sample) | PGI-RC<br>(default) | PGI-RC<br>(GREML unc.) |
| --- | --- | --- | --- | --- | --- | --- | --- |
| **Between-family results** |  |  |  |  |  |  |  |
| Polygenic index | 0.579***<br>(0.005) | 0.450***<br>(0.005) | 0.583***<br>(0.005) | 0.652***<br>(0.006) | 0.625***<br>(0.006) | 0.728***<br>(0.006) | 0.728***<br>(0.014) |
| First-stage estimate |  |  |  | 0.622***<br>(0.004) | 0.586***<br>(0.004) |  |  |
| First-stage $F$-statistic | | | | 27,261.16 | 20,791.79 | | |
| Incremental $R^2$ | 33.5% | 20.3% | 34.0% | | | | |
| Family fixed effects | NO | NO | NO | NO | NO | NO | NO |
| $N$ | 35,282 | 35,282 | 35,282 | 35,282 | 35,282 | 35,282 | 35,282 |
| **Within-family results** |  |  |  |  |  |  |  |
| Polygenic index | 0.521***<br>(0.007) | 0.415***<br>(0.007) | 0.537***<br>(0.007) | 0.614***<br>(0.008) | 0.571***<br>(0.009) |  |  |
| First-stage estimate |  |  |  | 0.600***<br>(0.005) | 0.558***<br>(0.005) |  |  |
| First-stage $F$-statistic | | | | 15,735.36 | 11,149.50 | | |
| Incremental $R^2$ | 27.1% | 17.2% | 28.9% | | | | |
| Family fixed effects | YES | YES | YES | YES | YES |  |  |
| $N$ | 35,282 | 35,282 | 35,282 | 35,282 | 35,282 | | |

**Table 4.**
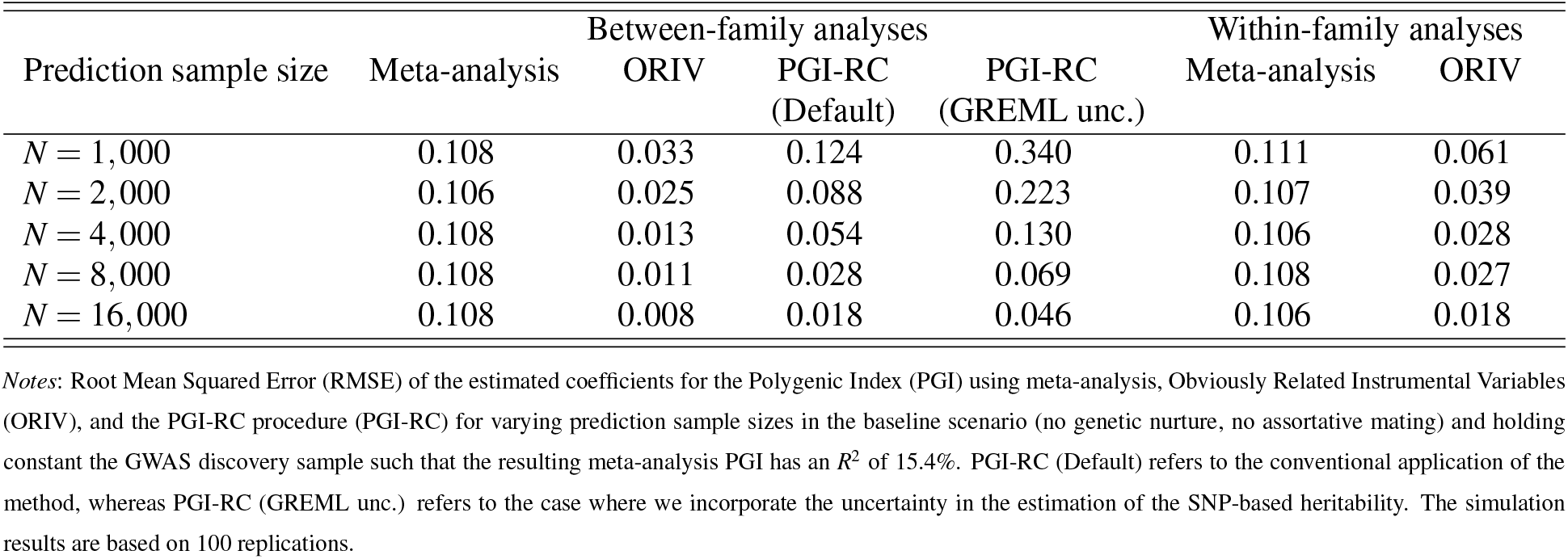
RMSE results accompanying Figure 1

**Table 5.**
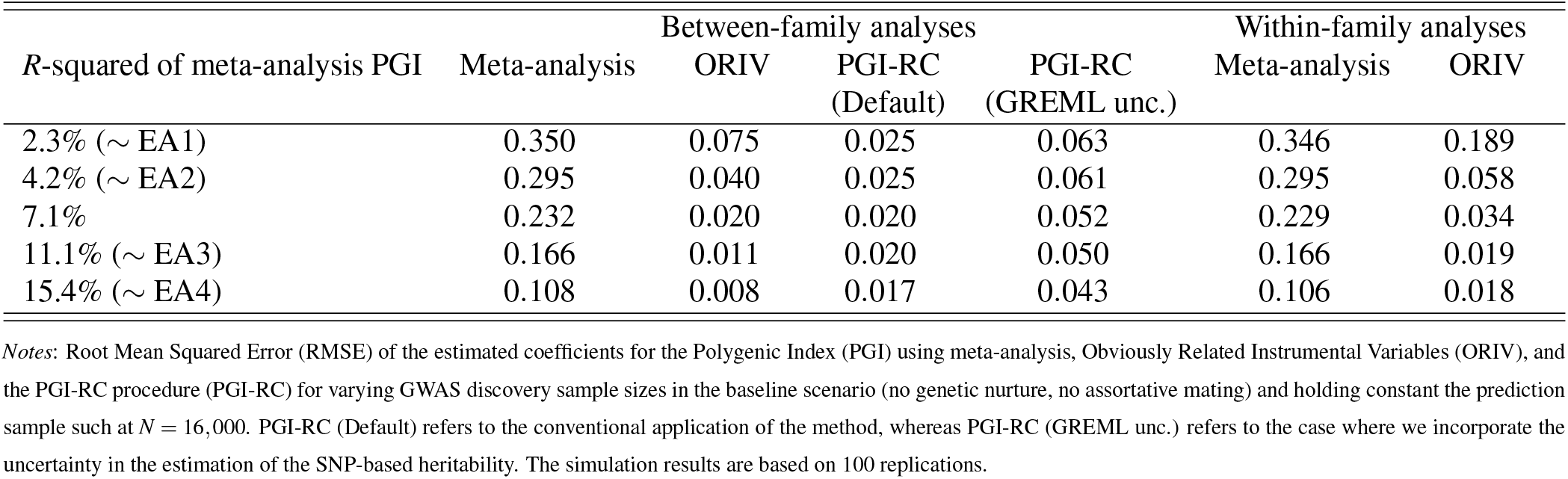
RMSE results accompanying Figure 2.

ORIV is somewhat biased when both the GWAS sample and the prediction sample are relatively small. This is not surprising, since it is well-known that the bias of IV is inversely proportional to the first-stage *F*-statistic (37; 38). As we derive in the Methods section, in this context, the first stage *F*-statistic is given by (see equation 24 for a derivation):

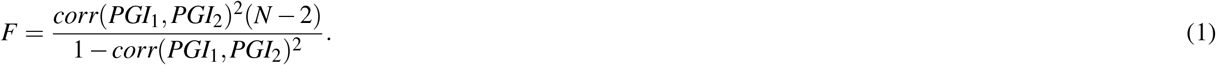

Thus, the bias in ORIV is determined by the correlation between the two PGIs, as well as the prediction sample size *N*. The correlation between the two PGIs is determined directly by the *R*-squared of the independent PGIs (see equations 9 and 16 in the Methods section). Therefore, with a small GWAS sample size *and* a small prediction sample size, ORIV is biased. Interestingly, when either the GWAS sample size increases (i.e., moving from EA2 to EA4 while holding constant the prediction sample at *N* = 1, 000) or the prediction sample size increases (i.e., moving from *N* = 1, 000 to *N* = 4, 000 or *N* = 16, 000 while holding the GWAS discovery sample constant), ORIV quickly converges to the true point estimate.

In this baseline scenario, the point estimates of the PGI-RC are largely independent of the GWAS sample size, but are very sensitive to a small prediction sample. The picture that emerges from Table 1 is that the PGI-RC outperforms ORIV when the prediction sample size is relatively large but the GWAS sample size is small; whereas ORIV tends to outperform the PGI-RC when the GWAS sample size is large but the prediction sample is relatively small. In case both the GWAS and prediction samples are small, both methods tend to estimate biased coefficients surrounded by wide confidence intervals, although the RMSE is still comfortably below that of a meta-analysis PGI (see Table 6 in Supplementary Information B).

**Table 6.** RMSE results accompanying Table 1.

| GWAS | Prediction sample | Meta-analysis | ORIV | PGI-RC<br>(Default) | PGI-RC<br>(GREML unc.) |
| --- | --- | --- | --- | --- | --- |
| ~EA2 | $N = 1,000$ | 0.291 | 0.106 | 0.116 | 0.334 |
| ~EA2 | $N = 4,000$ | 0.295 | 0.060 | 0.072 | 0.174 |
| ~EA2 | $N = 16,000$ | 0.296 | 0.035 | 0.023 | 0.058 |
| ~EA4 | $N = 1,000$ | 0.108 | 0.029 | 0.142 | 0.387 |
| ~EA4 | $N = 4,000$ | 0.114 | 0.013 | 0.046 | 0.118 |
| ~EA4 | $N = 16,000$ | 0.113 | 0.008 | 0.017 | 0.044 |

*Notes:* Root Mean Squared Error (RMSE) of the estimated coefficients for the Polygenic Index (PGI) using meta-analysis, Obviously Related Instrumental Variables (ORIV), and the PGI-RC procedure (PGI-RC) for the baseline scenario (no genetic nurture, no assortative mating), varying both the GWAS discovery sample and prediction sample. PGI-RC (Default) refers to the conventional application of the method, whereas PGI-RC (GREML unc.) refers to the case where we incorporate the uncertainty in the estimation of the SNP-based heritability. Between-family. The simulation results are based on 100 replications.

#### Genetic nurture

When genetic nurture is present, the results (Figure 3 and Table 7 in Supplementary Information B) very much resemble those obtained in the baseline case (Figure 1 and Figure 2). The coefficient of a meta-analysis PGI is consistently attenuated; ORIV and the PGI-RC both accurately target the correct point estimate, but confidence intervals are wide for the PGI-RC in small prediction samples, and for ORIV in small discovery samples. The most important difference is that with genetic nurture, the between- and within-family results are starting to diverge. In within-family designs, ORIV continues to outperform a meta-analysis PGI, yet slightly underestimates the true coefficient because of the noise introduced in the PGI since the GWAS did *not* control for genetic nurture (see Methods for a more extensive discussion).

**Figure 3.**
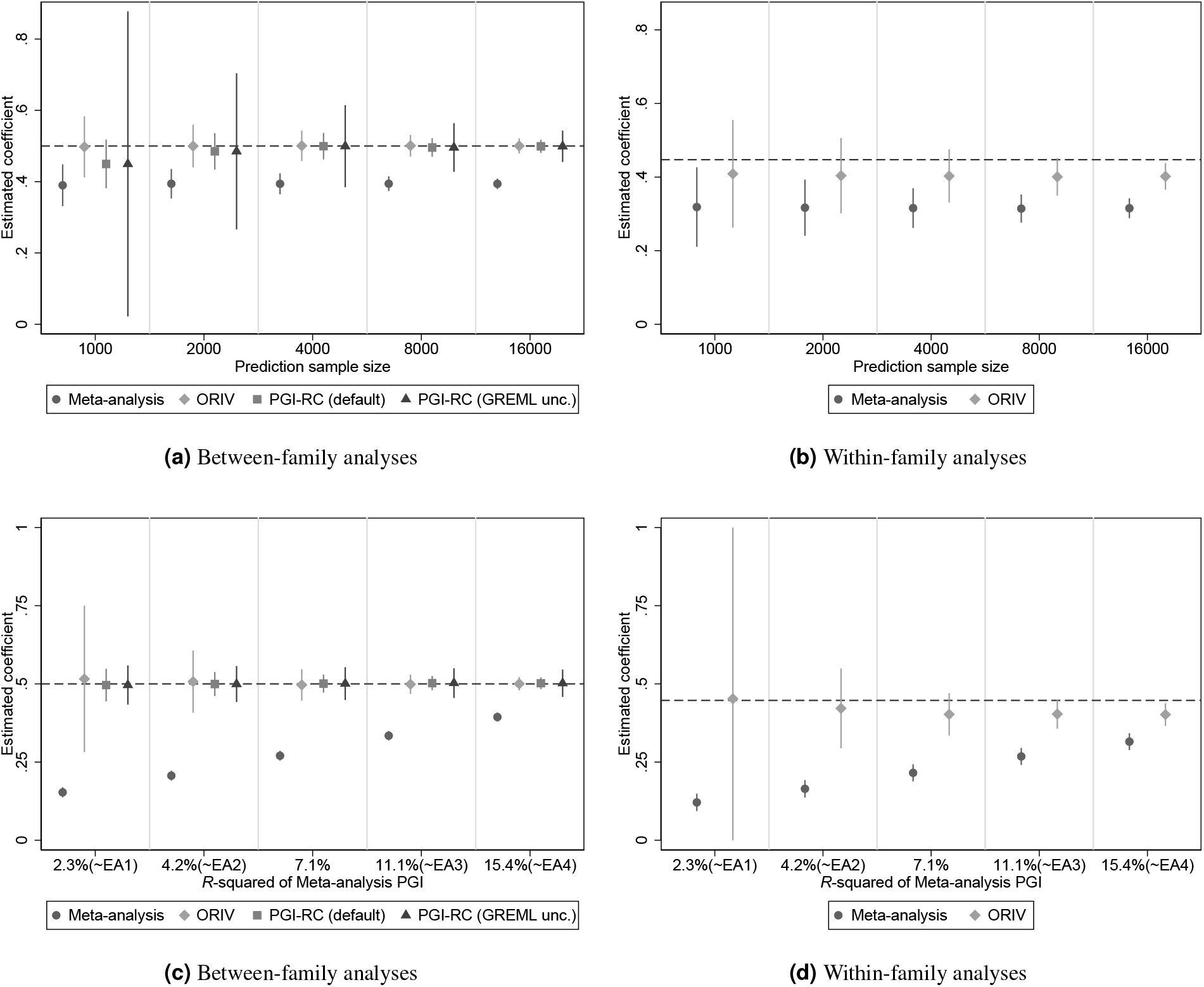
Estimated coefficients for the Polygenic Index (PGI) and their corresponding 95% confidence intervals using meta-analysis (circles), Obviously Related Instrumental Variables (ORIV, rhombuses), the default PGI-RC procedure (squares), and the PGI-RC procedure taking into account uncertainty in the GREML estimates (triangles). Top panels are for a scenario with genetic nurture but no assortative mating, and holding constant the GWAS discovery sample such that the resulting meta-analysis PGI has an *R*^2^ of 15.4%. Bottom panel is the same but now holding the discovery sample fixed at *N* = 16, 000. The dashed line represents the true coefficient. The simulation results are based on 100 replications.

**Table 7.** RMSE results accompanying Figure 3.

|  | Between-family analyses |  |  |  | Within-family analyses |  |
| --- | --- | --- | --- | --- | --- | --- |
|  | Meta-analysis | ORIV | PGI-RC<br>(Default) | PGI-RC<br>(GREML unc.) | Meta-analysis | ORIV |
| **Prediction sample size** |  |  |  |  |  |  |
| $N = 1,000$ | 0.105 | 0.029 | 0.125 | 0.344 | 0.129 | 0.060 |
| $N = 2,000$ | 0.109 | 0.024 | 0.087 | 0.223 | 0.130 | 0.054 |
| $N = 4,000$ | 0.105 | 0.016 | 0.055 | 0.131 | 0.131 | 0.048 |
| $N = 8,000$ | 0.107 | 0.012 | 0.028 | 0.070 | 0.133 | 0.048 |
| $N = 16,000$ | 0.106 | 0.007 | 0.018 | 0.046 | 0.132 | 0.046 |
| **R-squared of meta-analysis PGI** |  |  |  |  |  |  |
| 2.3% (~ EA1) | 0.347 | 0.081 | 0.021 | 0.057 | 0.326 | 0.187 |
| 4.2% (~ EA2) | 0.293 | 0.039 | 0.023 | 0.058 | 0.283 | 0.076 |
| 7.1% | 0.230 | 0.019 | 0.022 | 0.054 | 0.232 | 0.049 |
| 11.1% (~ EA3) | 0.166 | 0.011 | 0.020 | 0.049 | 0.179 | 0.044 |
| 15.4% (~ EA4) | 0.106 | 0.007 | 0.017 | 0.043 | 0.132 | 0.046 |

*Notes:* Root Mean Squared Error (RMSE) of the estimated coefficients for the Polygenic Index (PGI) using meta-analysis, Obviously Related Instrumental Variables (ORIV), and the PGI-RC procedure (PGI-RC) for varying prediction sample sizes in a scenario with genetic nurture but no assortative mating, and holding constant the GWAS discovery sample such that the resulting meta-analysis PGI has an $R^2$ of 15.4%. PGI-RC (Default) refers to the conventional application of the method, whereas PGI-RC (GREML unc.) refers to the case where we incorporate the uncertainty in the estimation of the SNP-based heritability. The simulation results are based on 100 replications.

#### Assortative mating

Figure 4 shows the results when in addition to genetic nurture, we add varying levels of assortative mating. All three methods are sensitive to assortative mating in between-family analyses (panel 4a), yet the PGI-RC procedure is affected more than ORIV (see also Table 8 in Supplementary Information B). This is not surprising, as the PGI-RC relies on an estimate of the SNP-based heritability obtained using GREML, which is known to be biased when assortative mating is present (39; 40). As expected, performing a within-family analysis (panel 4b) largely purges the bias induced by genetic nurture and assortative mating, with again ORIV providing a conservative estimate of the direct genetic effect in within-family analyses.

**Figure 4.**
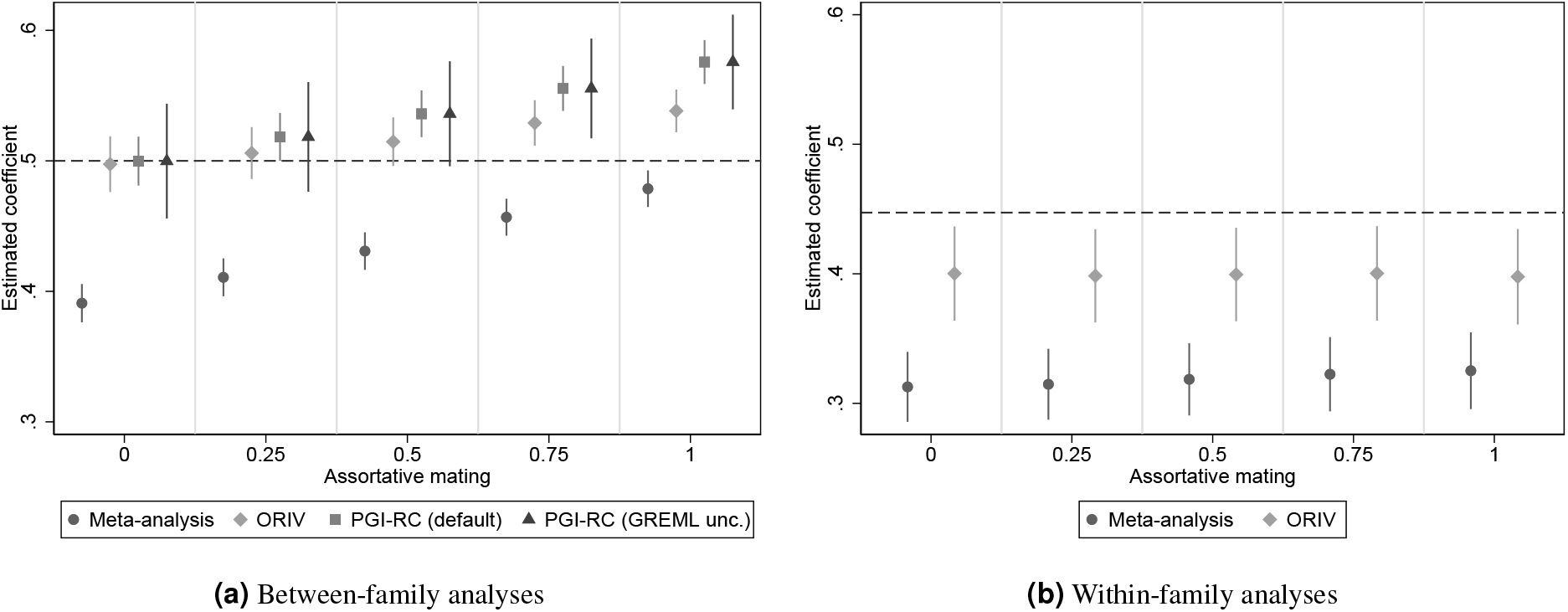
Estimated coefficients for the Polygenic Index (PGI) and their corresponding 95% confidence intervals using meta-analysis (circles), Obviously Related Instrumental Variables (ORIV, rhombuses), the default PGI-RC procedure (squares), and the PGI-RC procedure taking into account uncertainty in the GREML estimates (triangles) in a scenario with genetic nurture and varying levels of assortative mating (AM). The simulations hold constant the Genome-wide Association Study (GWAS) discovery sample such that the resulting meta-analysis PGI has an *R*^2^ of 15.4%, and the prediction sample size is held fixed at *N* = 16, 000. The dashed line represents the true coefficient. The simulation results are based on 100 replications.

**Table 8.**
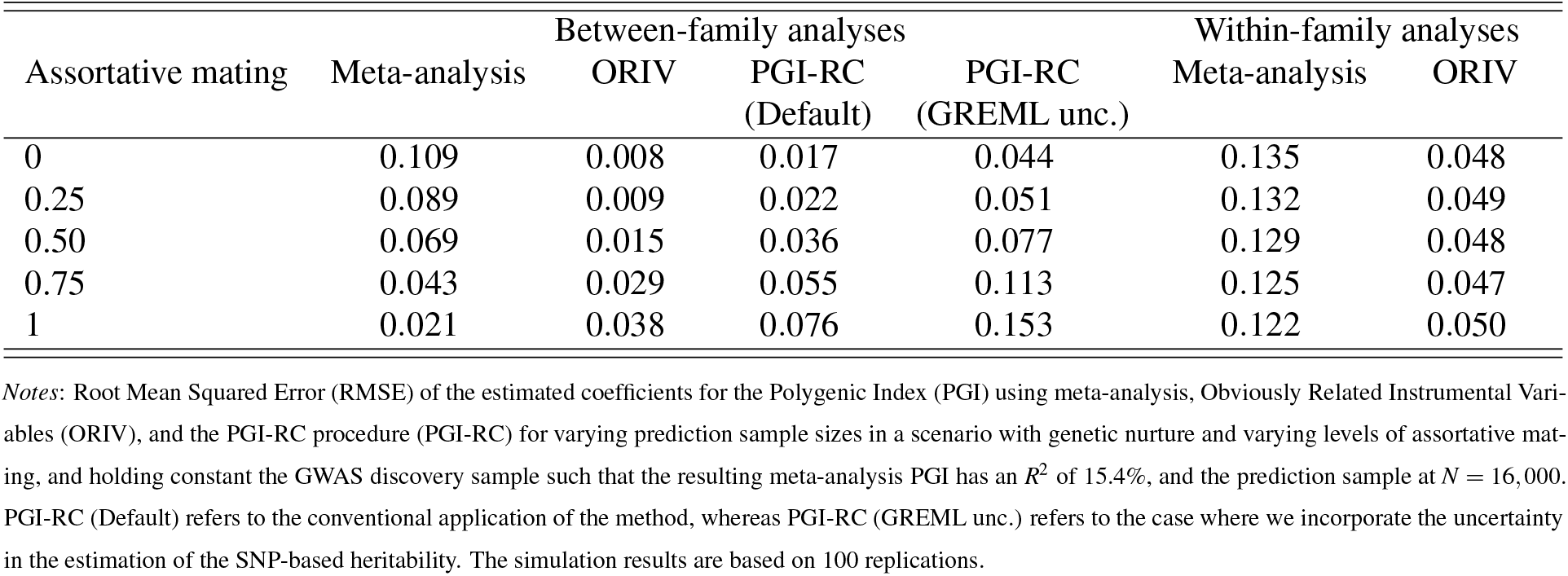
RMSE results accompanying Figure 4.

#### Imperfect genetic correlation

Figure 5 shows the sensitivity of the between-family approaches to an imperfect genetic correlation between the GWAS and prediction samples (left-panel) and between two GWAS samples that are meta-analysed or used for ORIV (right-panel). For a meta-analysis based PGI, these two settings are closely related because an imperfect genetic correlation between two GWAS samples by definition implies an imperfect genetic correlation with the prediction sample. The results show that both a meta-analysis PGI as well as ORIV are very sensitive to an imperfect genetic correlation between the discovery and prediction sample (see also Table 9 in Supplementary Information B for the corresponding RMSE values). In contrast, the PGI-RC procedure is not sensitive at all since it re-scales a particular coefficient with the SNP-based heritability *in the prediction sample*. For a lower genetic correlation across the two discovery samples, the PGI-RC is completely insensitive, and ORIV is remarkably robust against deviations from a perfect genetic correlation between the two GWAS discovery samples.

**Figure 5.**
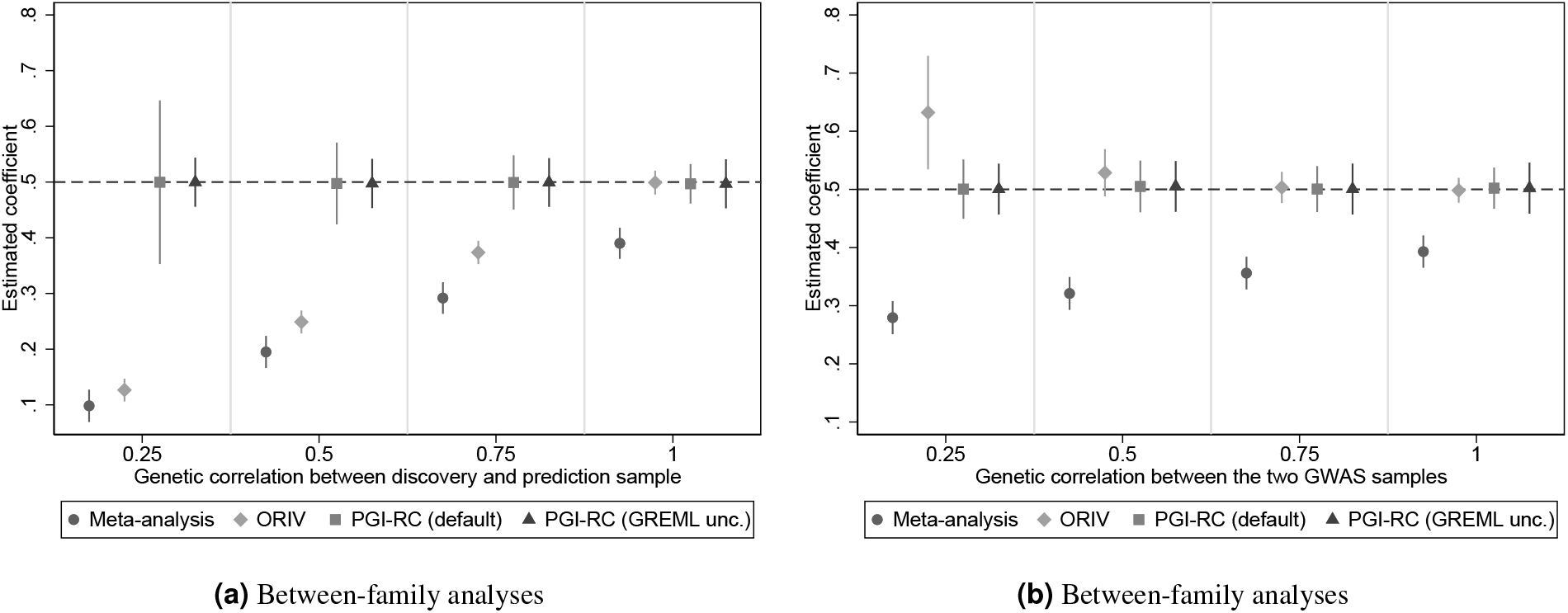
Estimated coefficients for the Polygenic Index (PGI) and their corresponding 95% confidence intervals using meta-analysis (circles), Obviously Related Instrumental Variables (ORIV, rhombuses), the default PGI-RC procedure (squares), and the PGI-RC procedure taking into account uncertainty in the GREML estimates (triangles) in a scenario without genetic nurture, without assortative mating, and varying levels of genetic correlation between the GWAS and prediction samples (left-panel) and between two GWAS samples that are meta-analysed or used by ORIV (right-panel). The simulations hold constant the Genome-wide Association Study (GWAS) discovery sample such that the resulting meta-analysis PGI has an *R*^2^ of 15.4%, and the prediction sample size is held fixed at *N* = 16, 000. Between-family analyses only. The dashed line represents the true coefficient. The simulation results are based on 100 replications.

**Table 9.** RMSE results accompanying Figure 5.

| $rG$ | $rG$ GWAS – prediction sample | | | | $rG$ two GWAS samples | | | |
| --- | --- | --- | --- | --- | --- | --- | --- | --- |
|  | Meta-analysis | ORIV | PGI-RC<br>(Default) | PGI-RC<br>(GREML unc.) | Meta-analysis | ORIV | PGI-RC<br>(Default) | PGI-RC<br>(GREML unc.) |
| 0.25 | 0.402 | 0.373 | 0.078 | 0.161 | 0.221 | 0.132 | 0.034 | 0.075 |
| 0.50 | 0.305 | 0.251 | 0.049 | 0.102 | 0.179 | 0.029 | 0.025 | 0.058 |
| 0.75 | 0.208 | 0.126 | 0.026 | 0.060 | 0.144 | 0.011 | 0.023 | 0.054 |
| 1 | 0.110 | 0.008 | 0.023 | 0.053 | 0.107 | 0.009 | 0.025 | 0.056 |

Notes: Root Mean Squared Error (RMSE) of the estimated coefficients for the Polygenic Index (PGI) using meta-analysis, Obviously Related Instrumental Variables (ORIV), and the PGI-RC procedure (PGI-RC) for varying levels of genetic correlation ( $rG$ ) in the baseline scenario without genetic nurture and no assortative mating, and holding constant the GWAS discovery sample such that the resulting meta-analysis PGI has an $R^2$ of 15.4% and the discovery sample fixed at $N = 16,000$ . PGI-RC (Default) refers to the conventional application of the method, whereas PGI-RC (GREML unc.) refers to the case where we incorporate the uncertainty in the estimation of the SNP-based heritability. The simulation results are based on 100 replications.

For a genetic correlation between the two GWAS samples of 0.75, ORIV is consistent, and even when the genetic correlation is as low as 0.5, the ORIV 95% confidence interval still includes the true coefficient.

#### Summary

In sum, the simulations suggest that it is virtually always beneficial to apply a measurement error correction compared to just using a meta-analysis based PGI. In choosing between measurement error corrections, PGI-RC tends to perform better than ORIV when the GWAS discovery sample is relatively small, or when there exists imperfect genetic correlation between the discovery and prediction sample. In contrast, the PGI-RC can exhibit considerable bias and uncertainty in case the prediction sample is small (*N <* 1, 000), and is biased for traits with considerable assortative mating. In those cases, as well as in within-family settings, ORIV seems the preferred alternative.

### Empirical illustration

In this section, we use OLS (using a meta-analysis based PGI), ORIV, and the PGI-RC to predict EA and height in a subsample of European ancestry siblings in the UK Biobank (*N* = 35, 282). In the prediction sample, we first residualized the outcomes EA and height for sex, year of birth, month of birth, sex interacted with year of birth, and the first 40 principal components of the genetic relationship matrix. For both EA and height, we consider three PGIs: (i) a PGI based on the UKB sample excluding siblings and their relatives; (ii) a PGI based on the 23andMe,Inc. sample (EA) or the GIANT consortium (height; (41)); and (iii) a PGI based on a meta-analysis of (i) and (ii). In addition, we construct two additional PGIs on the basis of randomly splitting the UKB discovery sample into two equal halves. All PGIs are constructed with the LDpred software (42) using a default prior value of 1. We standardize the PGIs to have mean 0 and standard deviation 1 in the analysis sample. The standardization of the PGI has the advantage that the square of its estimated coefficient in a univariate regression is equal to the *R*-squared (see Methods). Additionally, by standardizing a given PGI we can interpret the resulting coefficient as a one standard deviation increase in the true latent PGI (i.e., the additive SNP factor). More details on the variables and their construction can be found in Supplementary Information D.

#### SNP-based heritability

Using LDSC and GREML, we estimate the SNP-based heritability of EA in the prediction sample to be 0.162 (s.e. 0.027) and 0.155 (s.e. 0.019), respectively, which is somewhat lower than estimates from other samples in the literature. For height, the SNP-based heritability is estimated to be 0.497 (s.e. 0.039) using LDSC, and 0.530 (s.e. 0.020) using GREML. These results were obtained using one randomly selected sibling per family in the sibling subsample (*N* = 18, 989 for EA, *N* = 18, 913 for height). To obtain the LDSC estimate, the GWAS summary statistics were computed using FastGWA (43). In the GREML analysis (44), the analysis sample was slightly lower (*N* = 17, 696 for EA, *N* = 17, 849 for height) because we excluded closely related individuals using the default relatedness cut-off of 0.025. These SNP-based heritabilities are a useful benchmark, as they constitute an upper bound on the *R*^2^ we can achieve in our sample using a PGI (9). The SNP-based heritabilities additionally are a crucial input for disattenuating the OLS estimator in the PGI-RC (6). In fact, in the univariate case, the SNP-based heritability obtained using GREML *is* actually the PGI-RC estimate.

#### Educational attainment (EA)

Table 2 shows the results of regressions of residualized educational attainment (EA) in years (standardized to have mean 0 and standard deviation 1 in the sample) on the various PGIs. Figure 6a visualizes the results in terms of estimated heritability (i.e., the square of the standardized PGI coefficient). Meta-analyzing summary statistics from independent samples increases the standardized effect size and associated predictive power of the PGI compared with using the individual PGIs. That is, the PGI based on the meta-analysis of the UKB sample (excluding siblings and their relatives) and 23andMe delivers a standardized effect size of 0.276 (Column 3), implying an estimated heritability of 7.6%. This estimate is clearly higher than the effect sizes and heritability estimates obtained when using the UKB or 23andMe samples on their own (Columns 1 and 2) and the first two estimates of 6a. Nevertheless, the meta-analysis PGI still delivers an *R*^2^ that is substantially below the estimates of the SNP-based heritability of 15.5%-16.0%.^3^

**Figure 6.**
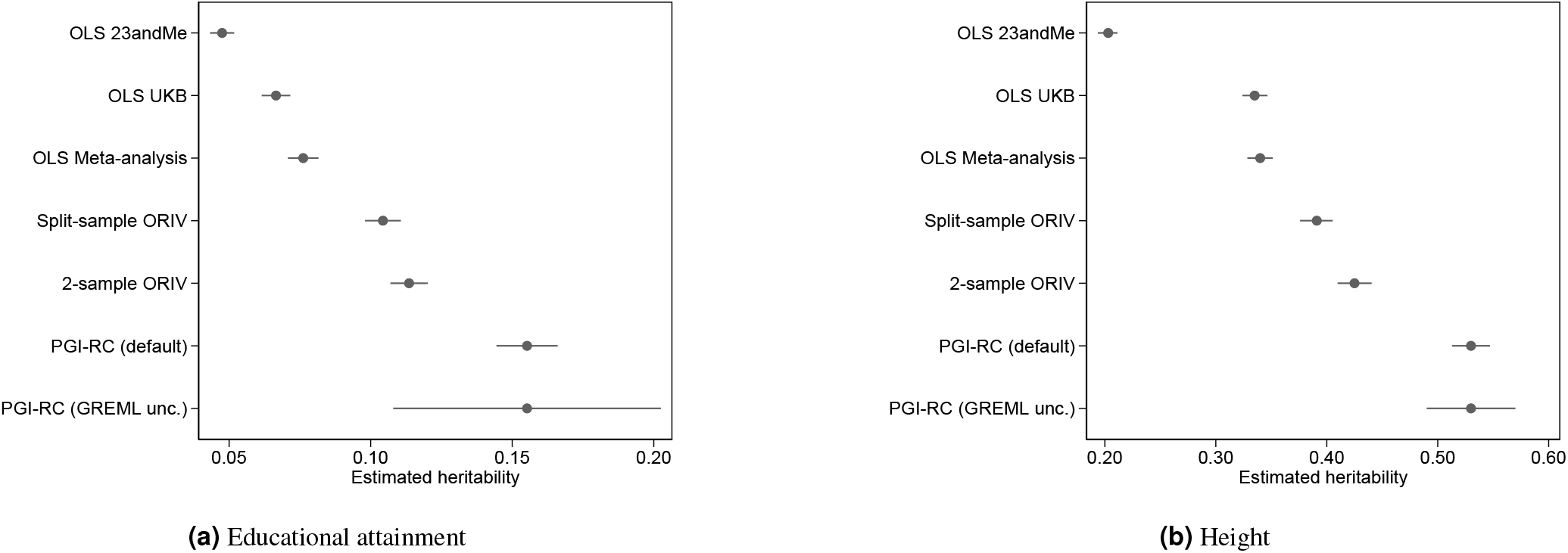
Results of the OLS and IV regressions in terms of implied heritability estimates and their 95% confidence intervals for **(a)** educational attainment (EA) and **(b)** height. The implied heritability is computed on the basis of the square of the standardized coefficients (see equation 23), and its standard error is obtained using the Delta method. Rest as in Table 2.

Column 4 of Table 2 shows the ORIV estimates employing the PGIs obtained from UKB and 23andMe as instrumental variables for each other. The ORIV standardized effect estimate is 0.337, which implies an estimated heritability 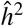 of 11.4%. While, unlike in the simulation, we do not know the true standardized effect size, it is reassuring that the implied heritability estimate is close to our empirical estimates of the SNP-based heritability of 15.5% (s.e. 1.9%; GREML) and 16.2% (s.e. 2.7%; LDSC), with the confidence intervals overlapping. In Column 5, we additionally present the ORIV results based on two PGIs that were constructed using two random halves of the UKB discovery sample. The IV assumptions are more likely to hold in this scenario since the samples are equally sized and they originate from the exact same environmental context. In particular, we estimate the genetic correlation between the 23andMe and UKB summary statistics to be 0.878 (s.e. 0.011), whereas the genetic correlation between the split-sample UKB summary statistics is 1.000 (s.e. <0.001). The resulting coefficient and implied heritability of 10.4% of the split-sample ORIV are only slightly below the two-sample ORIV results in Column 4, and considerably higher than the estimate obtained with the meta-analysis PGI.

When comparing ORIV to the PGI-RC, which in this univariate context equals the GREML estimate, we observe that the ORIV estimators are somewhat below the PGI-RC estimates (i.e., ORIV estimators may exhibit some bias). At the same time, when accounting for the uncertainty in the GREML estimate that underlies the PGI-RC (column 7 of Table 2 and bottom row of Figure 6a), the precision of ORIV estimators is considerably higher (i.e., ORIV estimators have lower variance). Hence, there is a bias-variance trade-off between ORIV and the PGI-RC in this context.

The results in the bottom panel of Table 2 are obtained using regressions that include family fixed effects. This approach only relies on within-family variation in the PGIs and therefore uncovers direct genetic effects (45). It bears repeating that in this context the PGI-RC cannot be applied, and so the relevant comparison here is between regular OLS and ORIV. The standardized effect estimates are substantially smaller within-families than they are between-families. This finding reflects an upward bias in the between-family estimates as a result of population phenomena, most notably genetic nurture (e.g., 15; 45; 46). More specifically, in line with the literature (e.g., 18; 47; 48), our within-family ORIV estimates are around 45% smaller than the between-family ORIV estimates. As in the between-family analyses, applying ORIV within-families increases the coefficient of the PGI compared to a meta-analysis or using standalone individual PGIs. ORIV estimates the standardized direct genetic effect to be 0.184 (using PGI from two samples – UKB and 23andMe) and 0.170 (using split-sample UKB PGIs). This estimate may still be prone to attenuation bias since the PGIs are based on a GWAS that did not consider indirect genetic effects from relatives (49; 50, see also Methods), but this estimate does represent a tighter lower bound on the direct genetic effect.

#### Height

Table 3 and Figure 6b present the results of the regressions with height as the outcome variable. The standardized effect sizes for height are considerably larger than for EA, consistent with the higher heritability of height. For example, a meta-analysis PGI based upon the UKB and the GIANT consortium GWAS summary statistics reaches a standardized effect size of 0.583, which corresponds to an incremental *R*-squared of 34%. It is also noteworthy that for height the between- and within-family results do not differ as much as they do for EA. Again, this is in line with the literature (e.g., 18) which generally finds genetic nurture to be more important for behavioral outcomes such as EA than for anthropometric outcomes like height.

Despite the differences in heritability and in the role of genetic nurture, we reach similar conclusions for height as for EA in the comparison of OLS (meta-analysis), ORIV, and the PGI-RC. The two-sample ORIV estimation is 25% (between-family) and even 30% (within-family) higher when using ORIV compared to a meta-analysis PGI. The confidence interval for the PGI-RC incorporating uncertainty in the GREML estimate is again larger than the confidence interval of the ORIV estimate, yet the point estimate of the PGI-RC is significantly higher than the one for ORIV. In the within-family analysis, ORIV delivers the tightest lower bound on the direct genetic effect for height, which is estimated to be above 0.6. With height being a typical trait to test new quantitative genetics methodologies (e.g., (44)), these empirical findings build confidence that our conclusions from the simulations apply more broadly.

#### Assortative mating

As shown in the simulations, depending on the degree of assortative mating, the SNP-based heritability may be overestimated, and hence the PGI-RC could be overestimating the true effect size more severely than ORIV. With EA being a trait that exhibits a considerable degree of assortative mating (51; 52; 53), it is not entirely clear whether the estimated SNP-based heritability of 15.5% (and corresponding standardized effect of ∼ 0.4) is the correct target, or that it is overestimating the true effect. Similarly, estimates of assortative mating for height are in the order of 0.23 (54), again leaving potential for bias in the GREML estimates that underlie the PGI-RC. For this reason, in Supplementary Information E.1, we compare the relative performance of ORIV and the PGI-RC for a trait with a similar level of heritability but that exhibits considerably less assortative mating: diastolic blood pressure (DBP). In line with the absence of assortative mating, the between-family and within-family results are highly concordant in case of DBP. Similar to the results for EA and height, ORIV increases the estimated coefficient by 27% compared to an OLS regression on basis of the meta-analyzed PGI. Again similar to EA and height, the confidence interval around the PGI-RC estimates is wider, but ORIV tends to produce slightly lower point estimates than the PGI-RC. In sum, it seems that our empirical findings are not driven by assortative mating in EA or height.

## Discussion

The increasing availability of genetic data over the last decade has stimulated genetic discovery in GWAS studies and has led to increases in the predictive power of polygenic indices (PGIs). Phenotypes such as educational attainment (EA) and height are currently at a critical turning point at which boosting the GWAS sample size further will only increase the predictive power of the PGIs at a marginal and diminishing rate. As a result, for the foreseeable future, regressions involving PGIs will be subject to an attenuation bias resulting from measurement error in the estimated GWAS coefficients. In this study, we compared two approaches that both attempt to overcome attenuation bias in the estimated regression coefficients of a PGI: Obviously-Related Instrumental Variables (ORIV) and the PGI Repository Correction (PGI-RC).

Extensive simulations show that in *between-family* analyses, the comparison of ORIV versus the PGI-RC (e.g., 6) is subject to a bias-variance trade-off that will differ across applications. The PGI-RC performs especially well compared with ORIV when there exists imperfect genetic correlation between the discovery and prediction sample, or when the GWAS discovery sample is relatively small. These conclusions hold even when incorporating the additional uncertainty induced by estimating the SNP-based heritability. However, when a sizable discovery sample is available that has near-perfect genetic correlation with the prediction sample, the RMSE of ORIV tends to be lower compared with the PGI-RC. Moreover, when there exists considerable assortative mating on the basis of the outcome variable, ORIV also tends to perform significantly better compared with the PGI-RC. In *within-family* analyses, ORIV is the most convenient way of estimating direct genetic effects, since the PGI-RC is not possible within-families. The simulations suggest that applying ORIV within-families will tighten the lower bound on the direct genetic effects in the presence of genetic nurture and assortative mating.

We empirically tested these predictions using UK Biobank data on educational attainment (EA) and height and largely confirm the simulation results. Both ORIV as well as the PGI-RC outperform a meta-analysis based PGI in terms of bias and RMSE. Among them, in our application, ORIV tends to underestimate the SNP-heritability somewhat (11% versus 15% for EA, and 43% versus 53% for height), but tends to have smaller standard errors than the PGI-RC, especially when the PGI-RC incorporates the uncertainty in the GREML estimates. On the basis of within-family analyses, in our application, ORIV estimated the direct genetic effect to be around 3.5% for EA and 38% for height, respectively a 30% (EA) and 14% (height) increase compared with a meta-analysis PGI.^4^ Similar findings for diastolic blood pressure (DBP) provide reassurance that the assortative mating on traits like EA and height were not driving these empirical findings.

There are alternative approaches to deal with measurement error than those analyzed in this study. Simulation-extrapolation (SIMEX, 55; 56) is an approach that also exploits external information on the SNP-based heritability, somewhat similar to the PGI-RC (6), yet relying on simulations. The advantage of ORIV over SIMEX is that it does not require external information or simulations, and can also be applied within-families. Notable other techniques to deal with measurement error are the Generalized Method of Moments (GMM, (57)) and Structural Equation Modeling (SEM, (25; 58)). Since IV can be seen as a special case of GMM and SEM models (e.g., 25; 59), the differences between the approaches are typically negligible in linear models. Although the distributional assumptions are somewhat stronger in SEM compared with IV, and including family fixed effects is possible (60; 61; 62) yet cumbersome, a possible advantage of SEM is its flexibility in allowing the factor loadings of the two individual PGIs to be different. This could be especially relevant when the sample sizes and/or genetic correlation with the prediction sample differ substantially across two GWAS discovery samples. An extensive comparison of ORIV versus (genomic) SEM or GMM is beyond the scope of this paper, but we anticipate that differences will typically be small unless the precision of the two independent PGIs differs substantially.

Whereas we have shown that both ORIV as well as the PGI-RC are superior to a regular meta-analysis PGI in terms of overcoming attentuation bias, this does not mean that further collection of additional genotyped samples is useless. In contrast, larger sample sizes are essential in identifying *specific* genetic variants that affect the phenotype of interest, allowing one to investigate the biological mechanisms driving these effects. Further, whereas the PGI-RC and ORIV are useful ‘scaling tools’ to estimate the regression coefficient of the latent ‘true’ PGI, they do not boost the predictive power of a PGI in out-of-sample applications. Finally, applying measurement error corrections using ORIV or the PGI-RC are not a substitute for within-family GWASs. The collection of family samples is the only way to explicitly control for the indirect genetic effects from relatives that plague the interpretation of the effects of PGIs in between-family studies. The collection of genetic data of family samples is on the rise (27), but their sample sizes are still comparatively small. The results of the present study suggest that the application of ORIV could help to reduce attenuation bias in PGIs constructed using the results of within-family GWASs.

## Methods

### Conceptual model

Consider a simple linear model in which a dependent variable *Y* (e.g., educational attainment) is influenced by many genetic variants:

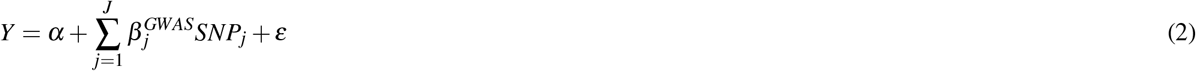

where *J* is the number of genetic variants (single-nucleotide polymorphisms, SNPs) included, *SNP*_*j*_ represents the number of effect alleles an individual possesses at locus *j*, and 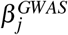 is the coefficient of SNP *j*. The true data generating process would also include the effects of maternal and paternal SNPs, because only conditional on parental genotypes the variation in SNPs is random and hence exogenous. We discuss the use of family data briefly below, but ignore the effects of parental genotype as well as environmental factors in the following discussion for simplicity.

The dependent variable *Y* is assumed to be standardized with mean zero and standard deviation 1 (*σ*_*Y*_ = 1). The true latent polygenic index *PGI*^*^ is then defined as:

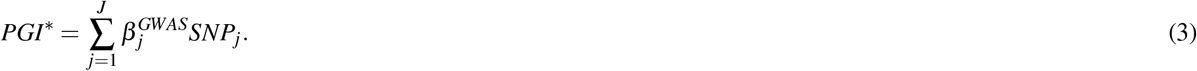

If we would observe the true polygenic index *PGI*^*^, then running the OLS regression

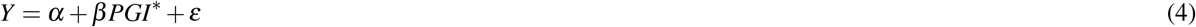

yields

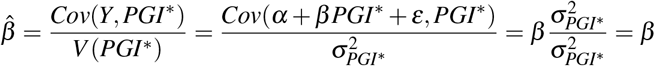

where *β* measures what happens to the outcome *Y* when the true latent *PGI*^*^ increases with 1 unit in the analysis sample. Since a 1 unit increase in the *PGI*^*^ is not straightforward to interpret, researchers are typically more interested in *β × σ*_*PGI*\*_, i.e., a one standard deviation increase in the true PGI. This estimate can be obtained by standardizing the PGI:

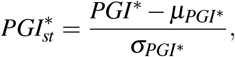

where *µ*_*PGI*\*_ is the mean of the true PGI, and *σ*_*PGI*\*_ is the standard deviation of the true PGI. If we now run the regression 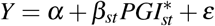, then the resulting estimator is:

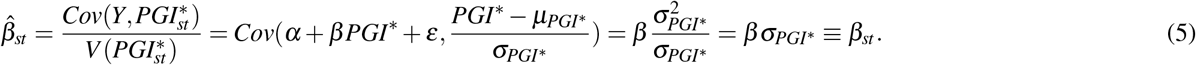

Apart from an arguably easier interpretation, the standardization of the PGI has the added advantage that there is a close connection between the estimated coefficient and the *R*-squared of this univariate regression. That is, in a univariate regression, the *R*-squared measures the squared correlation between the outcome and the independent variable, and it can be compared to the upper bound represented by the SNP-based heritability.

### Measurement error in the polygenic index

In practice, any estimated PGI is a proxy for the true latent polygenic index *PGI*^*^ because it is measured with error:

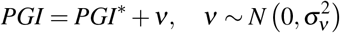

where we assume that the measurement error *v* is classical in the sense that it is uncorrelated to the error term in equation (2). If we estimate the regression *Y* = *α* + *β PGI* + *ε*, then measurement error in the *PGI* attenuates the coefficient of the *PGI* on *Y*:

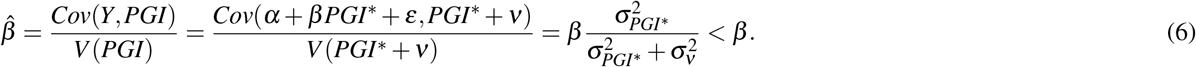

If – as is common in the literature – the observed PGI is standardized to obtain *PGI*_*st*_, it follows that:

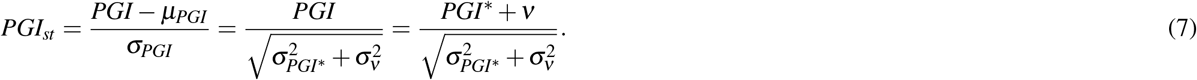

The resulting standardized coefficient of *PGI*_*st*_ on *Y* is given by

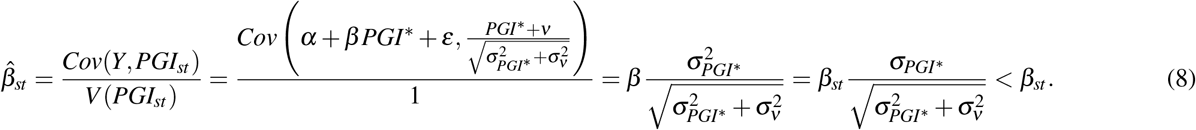

Note that standardizing the observed PGI with respect to its *own* standard deviation is a combination of standardizing with respect to the standard deviation of the true PGI as well as the standard deviation of measurement error (see also 25). Therefore, equation (8) shows that the estimate should be interpreted as the effect of a 1 standard deviation increase in the *observed* PGI (and not the true latent PGI). Hence, this estimate does not just underestimate the true *β* coefficient due to measurement error but should also be interpreted on a different scale than the effect of the true PGI (see e.g., 6; 25; 63).

Equation (8) can be rewritten as:

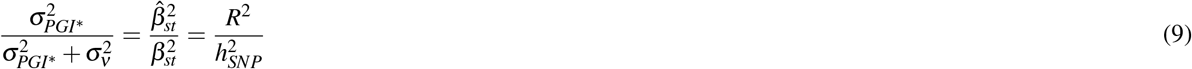

where we have used that the square of the standardized coefficient provides an estimate of the heritability. Hence, equation (9) shows that the bias in OLS is determined by the ratio of the estimated *R*-squared over the SNP-based heritability.

### PGI repository correction (PGI-RC)

Becker et al. (6) exploit their equivalent of equation (9) to directly derive an estimator that corrects for measurement error. The authors invert equation (9) and take the square root to obtain a disattenuation factor:^5^

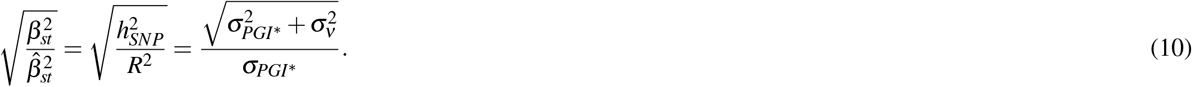

In turn, they suggest – in the univariate case – to multiply the estimated coefficient 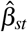 of the standardized PGI by the disattenuation factor to obtain effects of the true PGI (they call this the ‘the additive SNP factor’) free from measurement error:

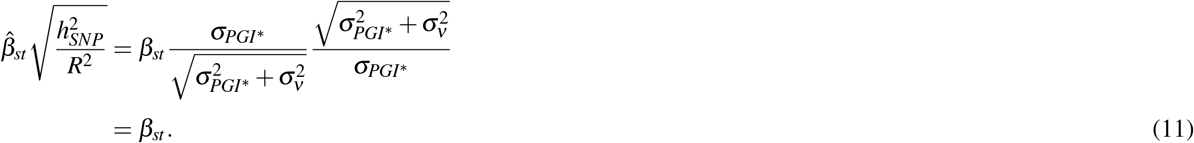

This procedure conveniently ensures that the estimated effect of the PGI is equal to the estimated SNP-based heritability, which the developers suggest to obtain using GREML. Hence, the bias of the product estimator in equation (11) is zero if the GREML assumptions are met and the researcher can compute the SNP-based heritability in the same sample as where the analysis is conducted.

The standard error of the PGI-RC estimator is discussed in the Supplementary Information of (6). When treating the factor 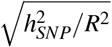 as a fixed non-stochastic scaling term, the variance can be derived using the Delta method:

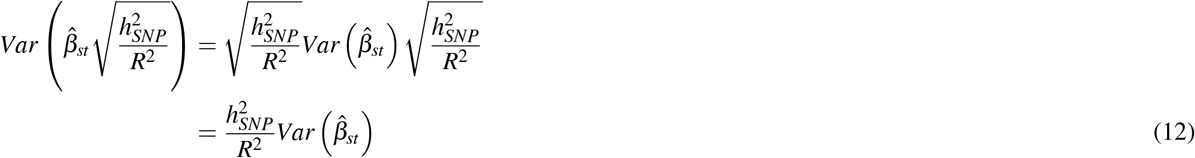

and the resulting standard error is then simply given by

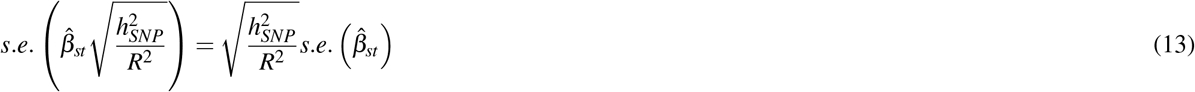

That is, both the coefficient as well as the standard error of the original standardized coefficient are simply scaled by the same factor 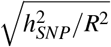. This standard error is treated as the default and also implemented in the accompanying software.

However, as acknowledged in (6, Supplementary Information, Section 5, Page 14-15), in practice the scaling factor is not a fixed immutable statistic, but rather a stochastic factor that is estimated in a first step. Hence, the true standard error of the PGI-RC equals the standard error of the square root of the SNP-based heritability obtained through GREML. Again using the Delta method, this leads to a standard error^6^

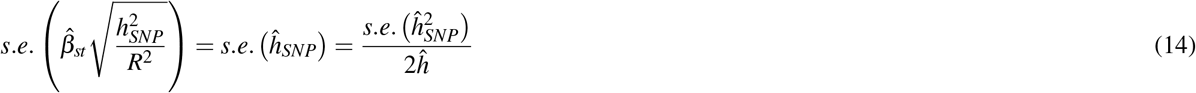

In our simulations as well as empirical applications, we report both standard errors (13) and (14) to allow for an accurate comparison across methods.

### Instrumental Variables

An alternative way of addressing measurement error is Instrumental Variables (IV regression). It has long been recognized in the econometrics literature (64; 65) that when at least two independent measures of the same construct (independent variable) are available, it is possible to retrieve a consistent effect of this construct on an outcome through IV estimation.^7^

In terms of formulas, if we have two measures for the true *PGI*^*^, *PGI*_1_ = *PGI*^*^ + *v*_1_ and *PGI*_2_ = *PGI*^*^ + *v*_2_, with *Cov*(*v*_1_, *v*_2_) = 0 and 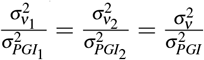. Then:

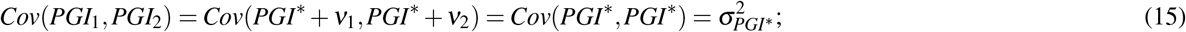

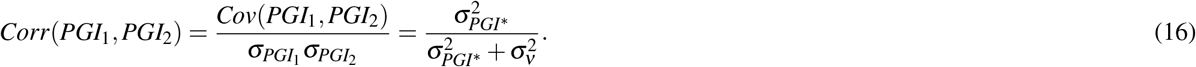

Hence, the correlation between the two PGIs can be used to correct for the attenuation bias that plagues the interpretation of the OLS estimates in equation (6). More formally, if we use *PGI*_2_ as an instrumental variable (IV) for *PGI*_1_, then the IV estimator is the ratio of the reduced form (regression of *Y* on *PGI*_2_) and the first stage (regression on *PGI*_1_ on *PGI*_2_):

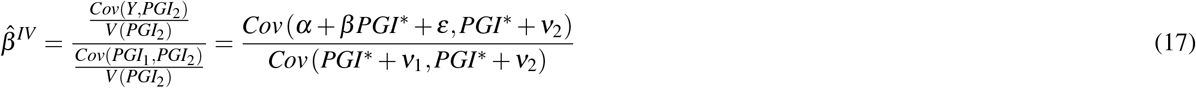

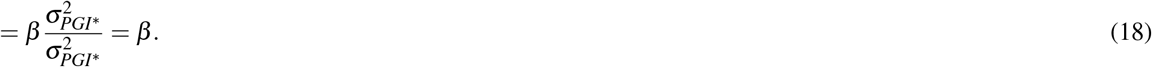

As a consequence, IV regression is able to estimate the unstandardized coefficient of the true latent PGI in a consistent way. But since unstandardized coefficients are hard to interpret, one is typically more interested in the standardized coefficient. In this case, the IV estimator is given by:

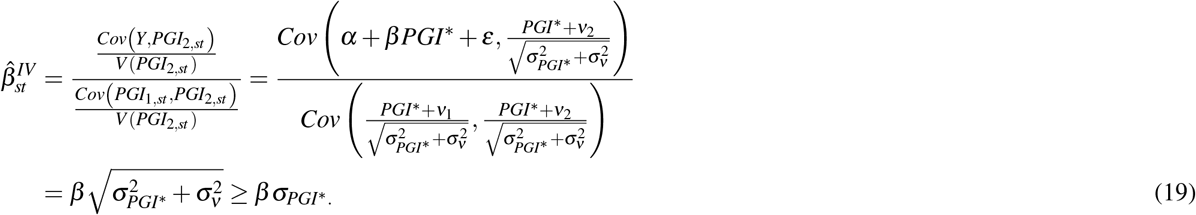

Importantly, the standardized IV estimator is not equal to the effect of a one standard deviation increase in the true PGI since the PGI is standardized with respect to the standard deviation of the observed instead of the true latent PGI. As a result, IV overestimates the true standardized coefficient. However, a way to retrieve the effect of a 1 standard deviation increase in the true latent PGI would be to scale the standardized IV coefficient:

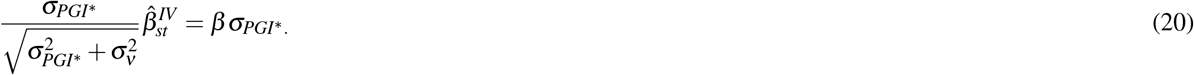

Although the scaling factor is unobserved, an estimate is given by the square root of the correlation between the two PGIs (see equation 16). Alternatively, one could also divide the observed standardized polygenic indices *PGI*_1,*st*_ and *PGI*_2,*st*_ by the same scaling factor:^8^

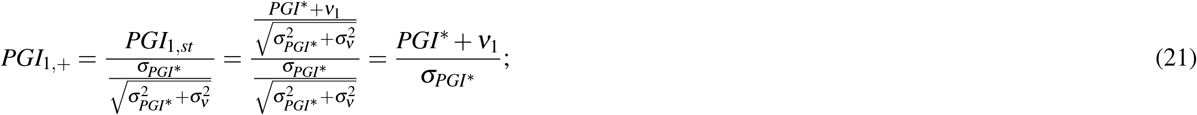

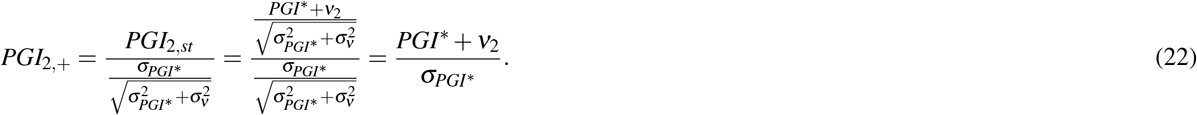

If we then base the IV estimator upon these scaled polygenic indices *PGI*_1,+_ and *PGI*_2,+_, then the resulting estimator is given by

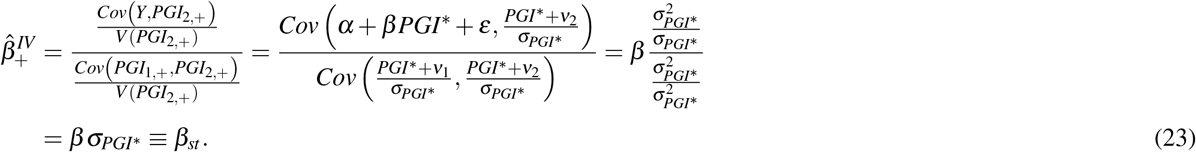

In sum, an IV estimate of the true standardized effect size can be obtained by (i) dividing the two independent standardized PGIs by the square root of their correlation,^9^ and (ii) using these scaled PGIs as instrumental variables for each other. This newly derived scaling factor avoids having to rescale regression estimates *ex post* to retrieve the estimated heritability as is done for example in (26).

#### Bias in IV

Instrumental Variable regression provides consistent estimates, yet it is biased in small samples with the bias in IV regression inversely related to the first-stage *F*-statistic with a factor roughly equal to 1*/F* (37; 38). The *F*-statistic in a univariate regression is equal to the square of the *t*-statistic of the first-stage coefficient 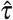. Therefore, in this context:

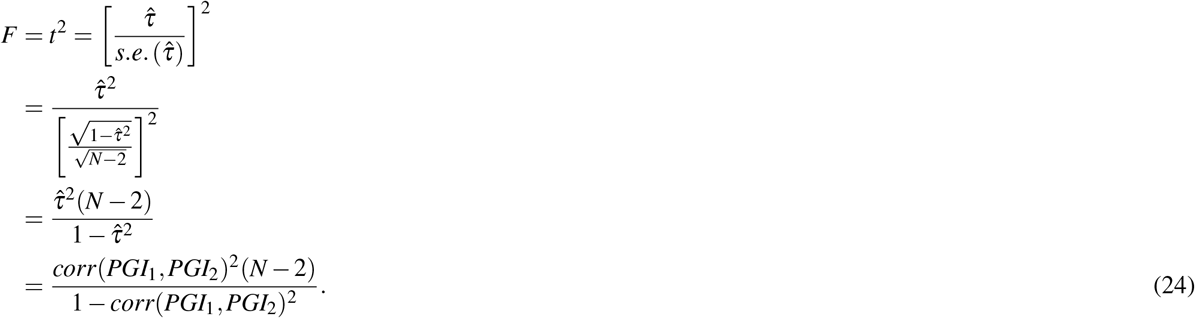

Where 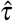 is the first-stage coefficient, and we have used the fact that the PGIs are standardized such that the coefficient 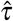 represents the correlation between the two PGIs. Hence, the performance of (OR)IV depends on the correlation between the two PGIs. This correlation is a function of the measurement error of the independent PGIs (see equation 16). Equation (24) also implies that, like OLS, the performance of (OR)IV is (largely) independent of the absolute values of *β* and 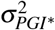. Unlike OLS, the bias of (OR)IV decreases with the prediction sample size *N*.

### Obviously-related Instrumental Variables

The most efficient implementation of the proposed IV estimator is the recently proposed technique “Obviously-Related Instrumental Variables” (ORIV) by (28). The idea is to use a ‘stacked’ model

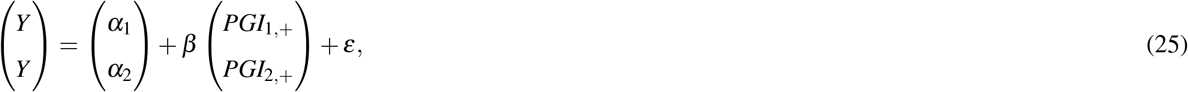

here one instruments the stack of estimated PGIs 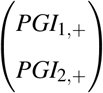 with the matrix

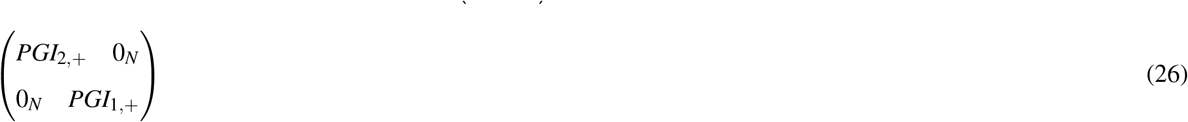

in which *N* is the number of individuals and *O*_*N*_ an *N ×* 1 vector with zero’s. The implementation of ORIV is straightforward: simply create a stacked dataset and run a Two-Stage Least Squares (2SLS) regression while clustering the standard errors at the individual level. In other words, replicate the dataset creating two values for each individual, and then generate two variables (i.e., an independent variable and an instrumental variable) that alternatively take the value of *PGI*_1,+_ and *PGI*_2,+_. The resulting estimate is the average of the estimates that one would get by instrumenting *PGI*_1,+_ by *PGI*_2,+_, and vice versa. This procedure makes most efficient use of the information in the two independent PGIs and avoids having to arbitrarily select one PGI as IV for the other. Family fixed effects can also be included in the model, in which case one should include a family-stack fixed effect in order to conduct only within-family comparisons within a stack of the data. Standard errors should then be clustered at both the family as well as the individual level. Supplementary Information E.3 illustrates the benefits of applying ORIV over regular IV in terms of point estimates, and a slight improvement in precision. Example syntax in Stata and R is available on our Github webpage https://github.com/geighei/ORIV.

### Within-family analysis

So far, we have ignored the potential influence of parental genotype on the individual’s outcome. Controlling for parental genotype is important since the genotype of the child is only truly random conditional on parental genotype. In other words, the only relevant omitted variables in a regression of an outcome on the child’s genotype are the genotype of the father and the mother. Leaving parental genotype out is not innocuous. As evidenced by several studies showing the difference between between-family and within-family analyses (e.g., 18), the role of parental genotype can be profound. Another way of showing this is by studying the effect of non-transmitted alleles of parents on their children’s outcomes (e.g., 47), to estimate so-called genetic nurture. The true data generating process (DGP) may therefore be:

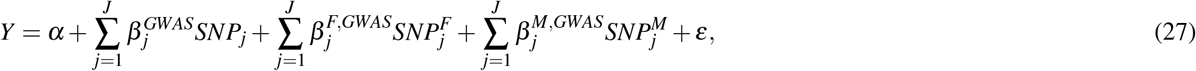

where the superscripts *F* and *M* denote father and mother, respectively. When the true DGP is governed by equation (27), 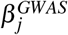 will be estimated with bias in case equation (2) is used in a GWAS. A simple solution would be to control for parental genotype or family fixed effects in the GWAS phase. However, with the recent exception of (27), GWAS discovery samples with sufficient parent-child trios or siblings are typically not available. Hence, a researcher often has no option but to work with the ‘standard GWAS’ coefficients that are obtained with equation (2) and that produce a biased PGI. Given this empirical reality, it is also the approach we adopt in our simulations.

In a between-family design, the bias in the coefficient of the resulting PGI tends to be upward, as the coefficients of the individuals and his/her parents are typically of the same sign (49). However, interestingly, when the conventional PGI is used in a within-family design, the bias is downward. The intuition is that when a conventional GWAS (i.e., a GWAS that does not control for parental genotype) is used in its construction, a PGI reflects direct genetic effects as well as indirect genetic effects (e.g., genetic nurture) arising from parental genotype. When applying these PGIs within-families, some of the differences in the PGI across siblings therefore spuriously reflect the effects of parental genotype, whereas in fact their parental genotype is identical. Hence, genetic nurture can be seen as measurement error in the PGI when applied in within-family analyses, leading to an attenuation bias (49).

A final source of downward bias could stem from indirect genetic effects, sometimes referred to as social genetic effects (50; 68), e.g., arising from siblings. For example, consider a case with two siblings where there is a direct effect *γ*_*j*_ of one’s sibling’s SNP on the outcome of the other:

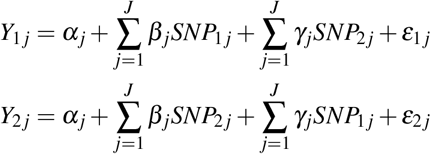

When taking sibling differences to eliminate the family fixed effects, we obtain:

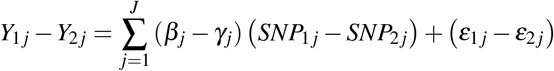

Since sibling effects are again likely to have the same sign as the direct effect, sibling effects cause a downward bias in the estimated effect of one’s own SNP, as measured by *β*_*j*_. Again, the only way to overcome this source of bias is to include the parental genotype in the GWAS, since conditional on the parental genotype, the genotypes of siblings are independent.

In sum, within-family analyses are the gold standard to estimate direct genetic effects, free from bias arising from the omission of parental genotype. However, when using a family fixed effects strategy on the basis of a PGI from a conventional GWAS that did not include parental genotype, the direct genetic effect is biased downward as a result of measurement error, genetic nurture effects and social genetic effects (see also 69). Therefore, this approach provides a lower bound estimate on the direct genetic effects.

## Acknowledgements

This research has been conducted using the UK Biobank Resource. The authors gratefully acknowledge participants of the 23andMe, Inc cohort for sharing GWAS summary statistics for educational attainment. This work made use of the Dutch national e-infrastructure with the support of the SURF Cooperative using grant no. EINF-2327. The authors also acknowledge funding from NORFACE through the Dynamic of Inequality across the Life Course (DIAL) programme (462-16-100), from the European Research Council (DONNI 851725 and GEPSI 946647), from the National Institute on Aging of the National Institutes of Health (RF1055654 and R56AG058726), and from the Dutch Research Council (016.VIDI.185.044). This research was supported by the National Institute for Health Research (NIHR Cambridge BRC-1215-20014 for EAWS). The views expressed are those of the authors and not necessarily those of the NIHR or the Department of Health and Social Care. We would like to thank Sjoerd van Alten, Dan Belsky, Neil Davies, Ben Domingue, Michel Nivard, and Elliot Tucker-Drob for valuable comments. The contents of this article are solely the responsibility of the authors and do not necessarily represent the official views of the National Institutes of Health.

## Supplementary Information

### A Simulations using GNAMES

In this section, we describe the outline of our simulations. For this purpose we developed a tool called GNAMES (Genetic Nurture and Assortative Mating Effects Simulator) available on https://github.com/devlaming/gnames. General usage and technical details (in terms of computational efficiency, indexing, etc.) are described in the GitHub repository.

#### A.1 Initialisation

We start with a population comprising *N* unrelated founders, for which we observe data on *M* independent SNPs. Let *G*_*ij*_ denote the genotype of founder *i* at SNP *j. G*_*ij*_ is drawn from Binom(2, *f*_*j*_), where *f*_*j*_ denotes the frequency of the coded allele for SNP *j*. Each allele frequency is sampled independently from a Beta(0.35, 0.35) distribution until it lies in [0.1, 0.9].

We draw an *M ×* 1 vector of direct SNP effects *β* and an *M ×* 1 vector of parental SNP effects *γ* (i.e., genetic nurture) as follows:

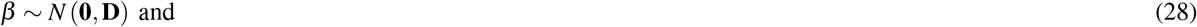

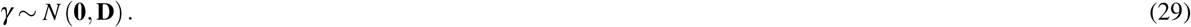

where *N*(·) denotes the Normal distribution, and **D** denotes a diagonal matrix, for which element *j, j* equals (2 *f*_*j*_ (1 − *f*_*j*_))^−1^. These SNP effects have the following properties:

- rarer variants have bigger effects than common variants, precisely to such an extent that a common and a rare SNP, on average, explain the same proportion of variation in the outcome, which aligns with SNP-effect assumptions made in, for instance, GREML analyses (44);
- for both the direct SNP effects and the genetic nurture effects, we assume complete polygenicity (i.e., all SNPs affect the outcome);
- genetic nurture and direct SNP effects are independent;
- SNP effects remain constant across generations (relative to each other; the overall scale may change to maintain target heritability).

#### A.2 Simulating phenotypes subject to genetic-nurture effects

In a given Generation *t*, comprising *N* individuals, let **G** denote the *N ×M* matrix of raw genotypes (i.e., count data between zero and two) and let **G**_*P*_ (resp. **G**_*M*_) denote the matrix of paternal (maternal) genotypes. The equations for the direct genetic component, the genetic nurture component, and the unique environment component can therefore be written as follows:

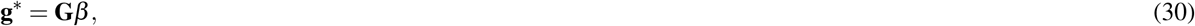

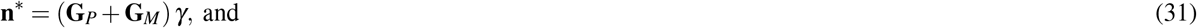

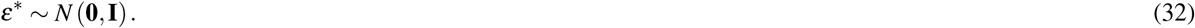

For the founders (i.e., Generation 0) we set **n**^*^ ∼ *N* (**0, I**).

To attain desired levels of SNP-based heritability (*h*^2^), the proportion of variance accounted for by genetic nurture (*n*^2^), and the unique environment, in each generation, we empirically standardise these components to have mean zero and unit variance, yielding **g, n**, and *ε*. Now, the equation for the outcome can be written as follows:

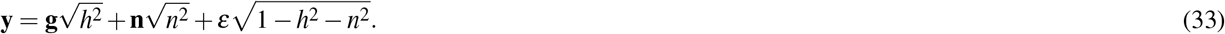

#### A.3 Assortative mating

In a given generation, we match pairs of individuals based on their value for the phenotype (i.e., we assign pairs of individuals to be mates). We follow the matching procedure as proposed by (40). More specifically, individuals are matched such that their phenotypic correlation equals the desired level *ρ*_*AM*_. This parameter is such that *ρ*_*AM*_ = 0 corresponds to random mating whereas *ρ*_*AM*_ = 1 correspond to assortative mating (AM) of such a degree that mates have a perfect phenotypic correlation.

We set up this procedure such that first-degree relatives in a given generation (i.e., siblings) cannot be matched. To this end, we assign Sibling 1 from each nuclear family to Group 1 and Sibling 2 from each nuclear family to Group 2, etc. Next, we apply the matching procedure by (40) within each group. The phenotypic correlation between mates then still equals the desired level *ρ*_*AM*_, even across groups (as borne out by our simulations).

Each mating pair has *C* = 2 children. Thus, if Generation *t* comprises *N* individuals (assuming *N* is an even number), there are *N/*2 mating pairs, yielding *CN/*2 = 2*N/*2 = *N* children in total. Hence, in Generation *t* + 1 there are also *N* individuals.

#### A.4 Partitioning data

We consider *T* = 10 generations. That is, we start at founders (*t* = 0), after which we have 10 generations of matching and mating. Assuming *N* is an even number, by the end of the simulation we have data on 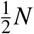 nuclear families. Per nuclear family, we have data on *C* = 2 siblings.

We partition the sibling data into three parts:

- GWAS Sample 1: Consider *N*_*GWAS*_ nuclear families, and per nuclear family consider only one of the two siblings for GWAS 1 (omitting parents).
- GWAS Sample 2: Consider *N*_*GWAS*_ nuclear families, and per nuclear family consider only one of the two siblings for GWAS 2 (omitting parents).
- Prediction Sample: Consider 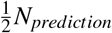 nuclear families, and per nuclear family consider both siblings (omitting parents), yielding a total prediction sample size of *N*_*prediction*_ (also assuming *N*_*prediction*_ is even).

These three samples are non-overlapping. By partitioning the data as such we can perform between-family GWASs and use the resulting SNP-effect estimates to construct PGIs that we can use for between- and within-family prediction. The sample size of both GWAS 1 and GWAS 2 is equal to *N*_*GWAS*_ here. The sample size of the meta-analysis equals *N*_*MA*_ = 2*N*_*GWAS*_. Finally, the sample size for prediction equals *N*_*prediction*_.

#### A.5 Imperfect genetic correlation

To investigate how imperfect genetic correlations between discovery and prediction samples impact the benchmark (OLS regression on the meta-analysis PGI), ORIV, and the PGI-RC, GNAMES also simulates a secondary trait **y**^*^ according to the following set of equations:

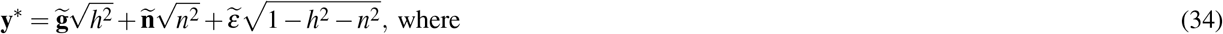

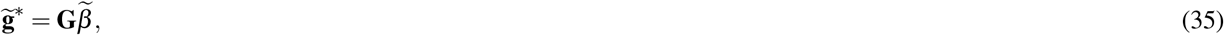

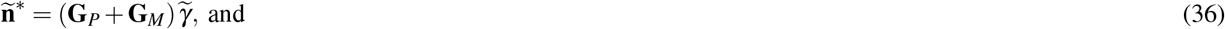

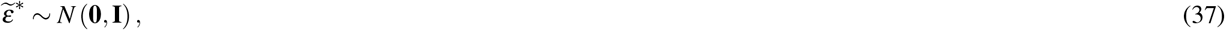

where 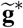 (resp. **ñ**^*^ and 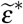) is standardised to have mean zero and unit variance, denoted by 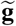 (**ñ** and 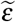). In these equations,

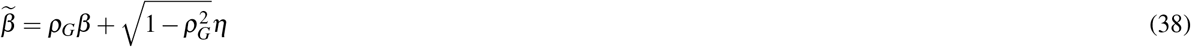

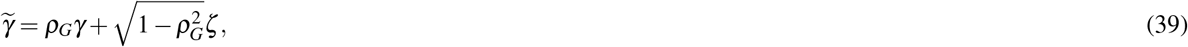

where

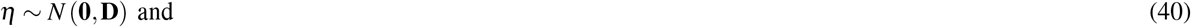

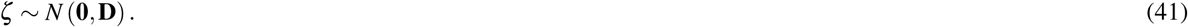

Under these definitions, in GWAS Sample 2 and in the Prediction Sample we may consider **y** while in GWAS Sample 1 we consider **y**^*^. In this case, the imperfect genetic correlation exists between one GWAS sample versus the other, while the prediction sample has a perfect genetic correlation with one of the two GWAS samples. Similarly, GWAS Sample 1 and GWAS Sample 2 may both consider **y** (i.e., both GWASs and the meta-analysis consider **y**), while the target trait in the prediction sample is **y**^*^. In this case, the imperfect genetic correlation exists only between prediction samples on the one hand, and GWAS samples on the other.

#### A.6 Simulation designs

We use various simulation designs to generate the results in the main paper. Below, we present each design with its input parameters. In addition to specifying parameter values, we also provide a link to the code for one run of that particular simulation design, and the corresponding Figure or Table in the paper. In a design where we fix *ρ*_*G*_ = 1, this is simply short-hand notation for using **y** both in GWAS 1, GWAS 2, and in the prediction sample (i.e., the secondary phenotype, **y**^*^ is ignored altogether in this case).

- *N*_*prediction*_ ∈ {1, 000; 2, 000; 4, 000; 8, 000; 16000} (Various prediction sample sizes)
  – For *M* = 5000, *h*^2^ = 20%, *n*^2^ = 10%, *ρ*_*AM*_ = 0, *ρ*_*G*_ = 1, and *N*_*GWAS*_ = 16000: see https://github.com/devlaming/gnames/blob/main/simulations/design3run1.py
  – This design is used in Figures 1 and 3.^10^
- *N*_*GWAS*_ ∈ {1, 000; 2, 000; 4, 000; 8, 000; 16000} (Various GWAS sample sizes)
  – For *M* = 5, 000, *h*^2^ = 25%, *n*^2^ = 0%, *ρ*_*AM*_ = 0, *ρ*_*G*_ = 1, and *N*_*prediction*_ = 16000: see https://github.com/devlaming/gnames/blob/main/simulations/design0run1.py
  – For *M* = 5, 000, *h*^2^ = 20%, *n*^2^ = 10%, *ρ*_*AM*_ = 0, *ρ*_*G*_ = 1, and *N*_*prediction*_ = 16000: see https://github.com/devlaming/gnames/blob/main/simulations/design1run1.py
  – These designs are used in Figure 2 and 3.
- *N*_*GWAS*_ ∈ {2, 000; 15, 000} and *N*_*prediction*_ ∈ {1, 000; 4, 000; 16, 000}
  – For *M* = 5000, *h*^2^ = 25%, *n*^2^ = 0%, *ρ*_*AM*_ = 0, *ρ*_*G*_ = 1: see https://github.com/devlaming/gnames/blob/main/simulations/design10run1.py
  – This design is used in Table 1.
- *ρ*_*AM*_ ∈ {0, 0.25, 0.5, 0.75, 1.0} (Various levels of AM)
  – For *M* = 5, 000, *h*^2^ = 20%, *n*^2^ = 10%, *ρ*_*G*_ = 1, *N*_*GWAS*_ = 16000, and *N*_*prediction*_ = 16, 000: see https://github.com/devlaming/gnames/blob/main/simulations/design2run1.py
  – This design is used in Figure 4.
- Perfect *ρ*_*G*_ between GWAS samples, but *ρ*_*G*_ ∈ {0, 0.25, 0.5, 0.75, 1} for GWAS versus prediction sample
  – For *M* = 5, 000, *h*^2^ = 25%, *n*^2^ = 0%, *N*_*GWAS*_ = 16, 000, and *N*_*prediction*_ = 16, 000: see https://github.com/devlaming/gnames/blob/main/simulations/design7run1.py
  – This design is used in Figure 5a.
- Perfect *ρ*_*G*_ between GWAS Sample 2 and Prediction Sample, but *ρ*_*G*_ ∈ {0, 0.25, 0.5, 0.75, 1} for GWAS Sample 1 versus GWAS Sample 2
  – For *M* = 5, 000, *h*^2^ = 25%, *n*^2^ = 0%, *N*_*GWAS*_ = 16, 000, and *N*_*prediction*_ = 16, 000: see https://github.com/devlaming/gnames/blob/main/simulations/design8run1.py
  – This design is used in Figure 5b.
- *N*_*GWAS*_ ∈ {3, 333; 25, 000} and *N*_*prediction*_ ∈ {1, 000; 4, 000; 16000}
  – For *M* = 5, 000, *h*^2^ = 15%, *n*^2^ = 0%, *ρ*_*AM*_ = 0, *ρ*_*G*_ = 1: see https://github.com/devlaming/gnames/blob/main/simulations/design9run1.py
  – This design is used in Table 10.
- *N*_*GWAS*_ ∈ {1, 000; 7, 500} and *N*_*prediction*_ ∈ {1, 000; 4, 000; 16000}
  – For *M* = 5, 000, *h*^2^ = 50%, *n*^2^ = 0%, *ρ*_*AM*_ = 0, *ρ*_*G*_ = 1: see https://github.com/devlaming/gnames/blob/main/simulations/design11run1.py
  – This design is used in Table 10.
- *ρ*_*AM*_ ∈ {0, 1} (Two levels of AM with high *M* and *N*_*GWAS*_)
  – For *M* = 100, 000, *h*^2^ = 20%, *n*^2^ = 10%, *ρ*_*G*_ = 1, *N*_*GWAS*_ = 50, 000, *N*_*prediction*_ = 16, 000: see https://github.com/devlaming/gnames/blob/main/simulations/design4run1.py
  – This design is used in Figure 8.

**Table 10.** Estimated coefficients for the Polygenic Index (PGI) and their corresponding 95% confidence intervals using meta-analysis, Obviously Related Instrumental Variables (ORIV), and the PGI-RC procedure in the baseline scenario (no genetic nurture, no assortative mating; between-family analyses only).

| GWAS | Prediction sample | Meta-analysis | ORIV | PGI-RC<br>(Default) | PGI-RC<br>(GREML unc.) |
| --- | --- | --- | --- | --- | --- |
| SNP-based heritability = 0.15 ($\beta = 0.387$) | | | | | |
| $R^2 / h_{SNP}^2 = 0.16$ | $N = 1,000$ | 0.160<br>(0.098 – 0.222) | 0.435<br>(-0.150 – 1.021) | 0.384<br>(0.231 – 0.537) | 0.384<br>(-0.054 – 0.823) |
| $R^2 / h_{SNP}^2 = 0.16$ | $N = 4,000$ | 0.159<br>(0.128 – 0.190) | 0.384<br>(0.229 – 0.538) | 0.379<br>(0.304 – 0.453) | 0.379<br>(0.191 – 0.566) |
| $R^2 / h_{SNP}^2 = 0.16$ | $N = 16,000$ | 0.158<br>(0.142 – 0.173) | 0.383<br>(0.306 – 0.459) | 0.389<br>(0.351 – 0.428) | 0.389<br>(0.320 – 0.458) |
| $R^2 / h_{SNP}^2 = 0.6$ | $N = 1,000$ | 0.303<br>(0.244 – 0.363) | 0.392<br>(0.307 – 0.478) | 0.327<br>(0.263 – 0.391) | 0.327<br>(-0.101 – 0.756) |
| $R^2 / h_{SNP}^2 = 0.6$ | $N = 4,000$ | 0.299<br>(0.269 – 0.329) | 0.386<br>(0.344 – 0.429) | 0.387<br>(0.349 – 0.426) | 0.387<br>(0.233 – 0.541) |
| $R^2 / h_{SNP}^2 = 0.6$ | $N = 16,000$ | 0.300<br>(0.285 – 0.315) | 0.387<br>(0.366 – 0.408) | 0.385<br>(0.366 – 0.405) | 0.385<br>(0.331 – 0.440) |
| SNP-based heritability = 0.5 ($\beta = 0.707$) | | | | | |
| $R^2 / h_{SNP}^2 = 0.16$ | $N = 1,000$ | 0.296<br>(0.234 – 0.358) | 0.722<br>(0.123 – 1.321) | 0.682<br>(0.536 – 0.827) | 0.682<br>(0.320 – 1.043) |
| $R^2 / h_{SNP}^2 = 0.16$ | $N = 4,000$ | 0.293<br>(0.262 – 0.324) | 0.700<br>(0.445 – 0.955) | 0.705<br>(0.630 – 0.781) | 0.705<br>(0.588 – 0.823) |
| $R^2 / h_{SNP}^2 = 0.16$ | $N = 16,000$ | 0.290<br>(0.274 – 0.305) | 0.695<br>(0.571 – 0.819) | 0.707<br>(0.669 – 0.745) | 0.707<br>(0.668 – 0.746) |
| $R^2 / h_{SNP}^2 = 0.6$ | $N = 1,000$ | 0.552<br>(0.499 – 0.605) | 0.707<br>(0.614 – 0.799) | 0.660<br>(0.596 – 0.723) | 0.660<br>(0.322 – 0.997) |
| $R^2 / h_{SNP}^2 = 0.6$ | $N = 4,000$ | 0.548<br>(0.521 – 0.575) | 0.705<br>(0.658 – 0.752) | 0.709<br>(0.675 – 0.744) | 0.709<br>(0.628 – 0.791) |
| $R^2 / h_{SNP}^2 = 0.6$ | $N = 16,000$ | 0.546<br>(0.533 – 0.560) | 0.702<br>(0.679 – 0.725) | 0.707<br>(0.689 – 0.724) | 0.707<br>(0.677 – 0.737) |

Notes: Coefficients of an OLS regression on a meta-analysis based PGI (column 3), a 2SLS regression using ORIV (column 4), the default PGI-RC (column 5), and the PGI-RC incorporating uncertainty of the GREML estimate (column 6). 95% confidence intervals in parentheses. The simulation results are based on 100 replications.

#### A.7 Meta-analysis as implemented in GNAMES

In our simulations using GNAMES, we pool the data used in the two main GWASs, and use this pooled data for a third GWAS, which we refer to as a meta-analysis. Although the latter technically is not a meta-analysis, we here show that this pooled simple GWAS is equivalent to a meta-analysis. As such, we can safely refer to the pooled GWAS as a meta-analysis within the context of our simulations.

We assume there are no control variables to take into account. Moreover, we assume both the SNP and outcome have been standardised to mean zero and unit variance, and we assume a given SNP individually to explain only a negligible proportion of the variation in the outcome. Consider SNP **x** and outcome **y** (both *N ×* 1 vectors). We split the sample in two chunks with size *N*_1_ and *N*_2_, such that *N* = *N*_1_ + *N*_2_. Let *i* = 1, 2 denote an index for the two corresponding subsamples.

The GWAS estimate, for the given SNP, in Subsample *i* = 1, 2 can be written as follows:

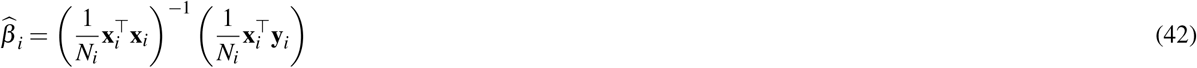

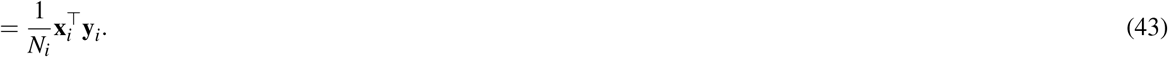

where **x**_*i*_ (resp. **y**_*i*_) denotes the SNP (outcome) vector in Subsample *i*, and where the last equality holds because SNPs have been standardised (i.e., 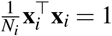)

Moreover, the GWAS *z*-score in Subsample *i* can be written as

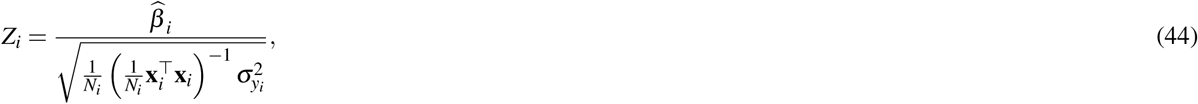

where 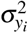 is the phenotypic variance in Subsample *i* not accounted for by the given SNP. As SNPs are standardised, we have that 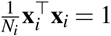. Moreover, as the SNP explains hardly any variation in *y*, we have that 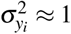. Therefore, we can approximate the right-hand side in Equation 44 as follows:

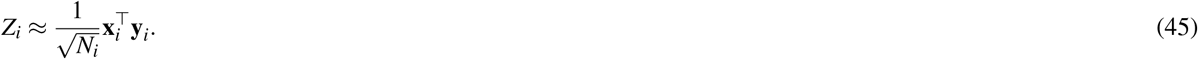

Moreover, under this standardisation the following approximation of the SNP-effect estimate as function of its *z*-score also holds:

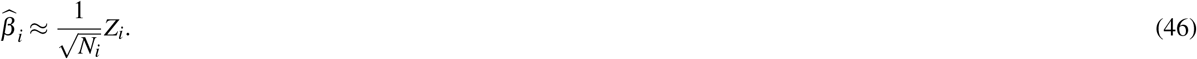

The sample-size weighted meta-analyses *z*-score (70) is given by

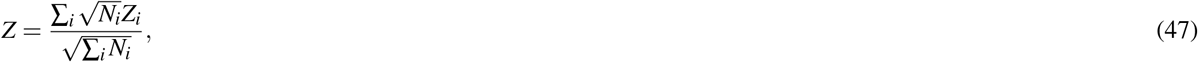

which for the case of two subsamples reduces to

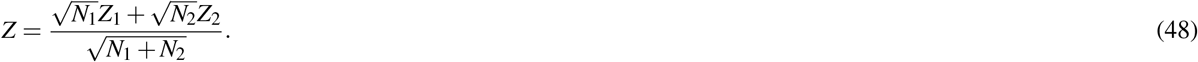

Using Equation 45, the meta-analysis *z*-score can be approximated as follows:

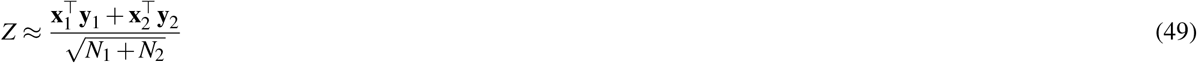

Moreover, using our approximation for 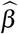 as a function of its *z*-score in Equation 46, we can approximate our meta-analysis SNP-effect estimate as a function of the meta-analysis *z*-score:

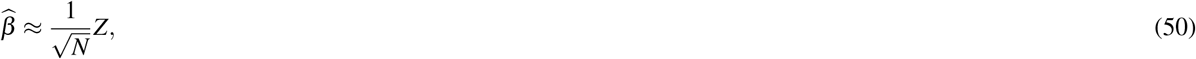

where *N* = *N*_1_ + *N*_2_. Combining Eq. 49 and 50 yields that the meta-analysis SNP-effect estimate can be approximated as follows:

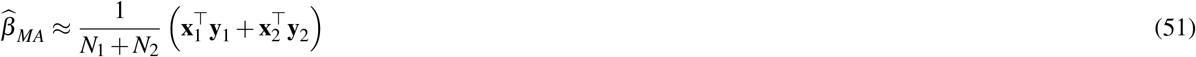

Now, consider the SNP-effect estimate by applying ordinary least squares (OLS) to pooled data directly (instead of using meta-analysis). This OLS estimate equals:

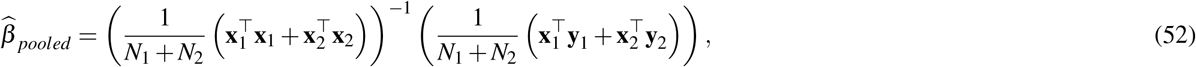

where 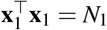 and 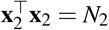 by virtue of aforementioned SNP standardisation. Thus, this expression reduces to

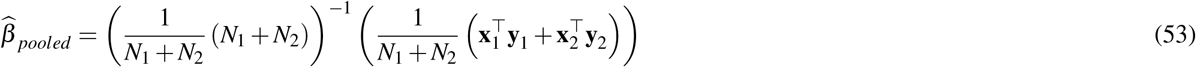

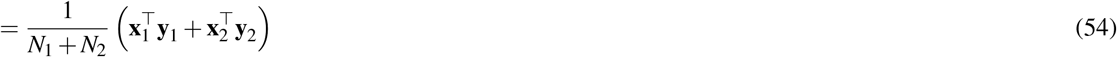

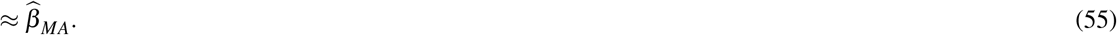

Similarly, we also have that *Z*_*pooled*_ ≈ *Z*_*MA*_. This finding aligns with the work by (71), showingthat there are no gains in statistical efficiency by analysing pooled data in a GWAS instead of using meta-analysis.

To illustrate the equivalence, we simulate SNP and outcome data, which we then split into two subsamples. We perform a GWAS in each subsample as well as in the pooled data. Finally, we meta-analyse results from the two subsamples, and compare these to results from the GWAS on pooled data. Figure 7 shows pooled GWAS *z*-scores versus meta-analysis *z*-scores. The two sets of *z*-scores are virtually indistinguishable; they have the same scale and an *R*^2^ of 99.99%. Therefore, in the context of GNAMES, we can talk about a pooled GWAS and meta-analysis interchangeably.

**Figure 7.**
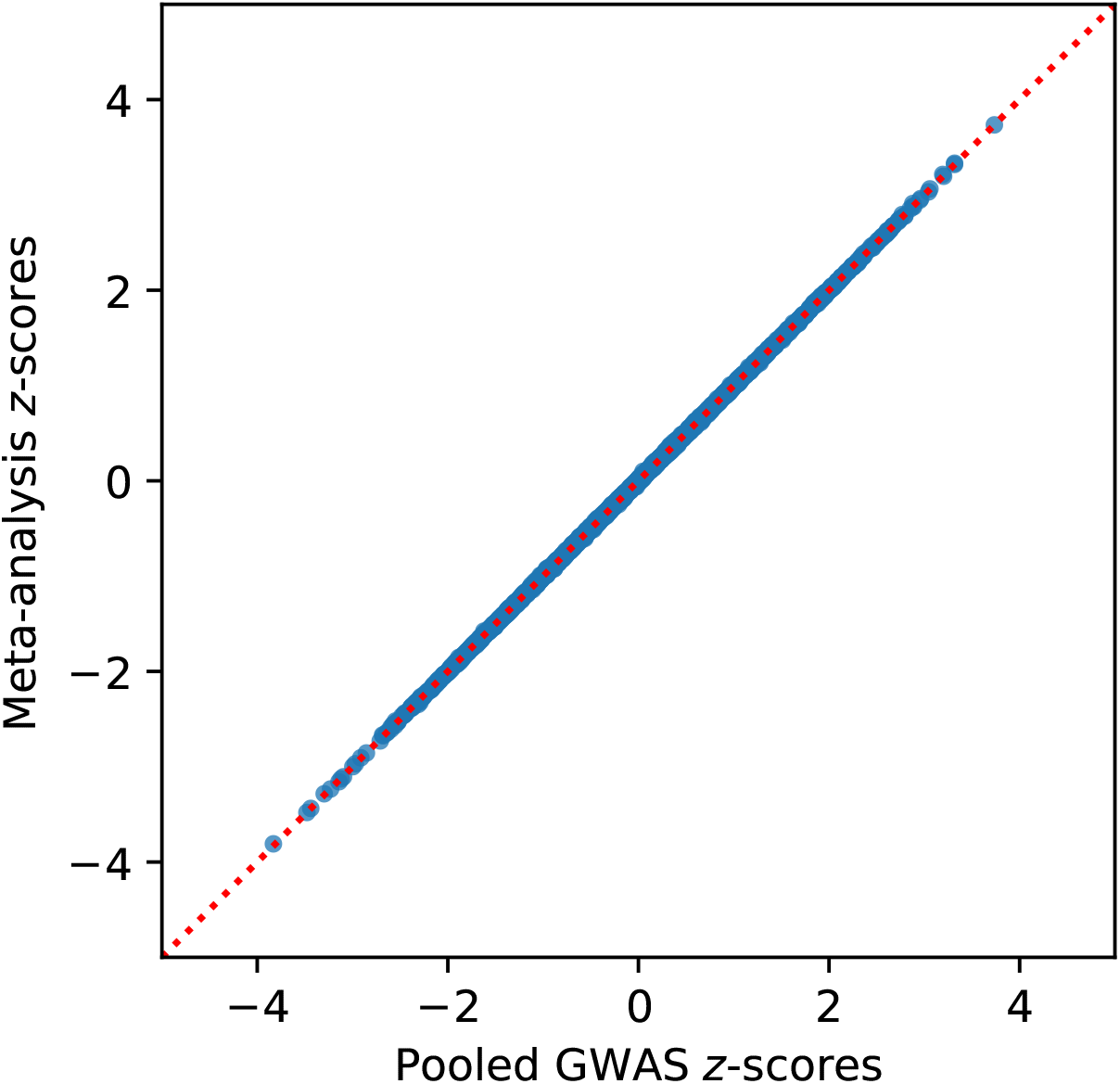
Pooled GWAS *z*-scores versus meta-analysis *z*-scores (*R*^2^ = 99.99%)

## B Root Mean Squared Error (RMSE)

In this section we present the Root Mean Squared Error (RMSE) corresponding to each of the figures in the main text.

## C Robustness simulation results

### C.1 Choice of 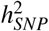

In our main set of simulations we set the SNP-based heritability to 25%, corresponding to *β*_*st*_ = 0.5. Here, we seek to investigate the generalizibility of our simulation findings to different levels of SNP-based heritability. In particular, we decrease the SNP-based heritability to 0.15 (i.e., roughly the SNP-based heritability of EA in our empirical analysis) and to 0.5 (i.e., roughly the SNP-based heritability of height in our empirical analysis). In investigating scenarios, for comparability, we maintain the same ratio of the *R*-squared (*R*^2^) to the SNP-based heritability 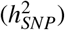 as we use in Table 1. That is, since EA2 roughly corresponds to a ratio of 0.16(= 0.042*/*0.25) and *EA*4 corresponds to a ratio of 0.6(= 0.15*/*0.25), we investigate a similar ratio for the other two levels of heritability. Table 10 presents the results.

Similar to Table 1, ORIV is somewhat biased for relatively small GWAS samples, while the PGI-RC is somewhat sensitive to a small prediction sample. Still, both methods clearly outperform a meta-analysis based PGI. For larger GWAS and prediction samples, the two methods both perform accurately, with little differences between them. Applying the default PGI-RC produces slightly narrower confidence intervals. Together, these simulation results show that the performance of a meta-analysis based PGI, ORIV or the PGI-RC are not driven by the absolute value of *β*_*st*_ or the SNP-based heritability, such that our conclusions are not specific to any level of SNP-based heritability.

### C.2 Sample size GWAS

The expected *R*-squared of a given PGI in a regression depends on the ratio of the number of SNPs over the GWAS discovery sample. Therefore, to keep the simulation space manageable we proportionally reduced both the number of SNPs as well as the GWAS discovery sample to generate realistic values for the *R*-squared of the PGIs. In order to verify that this shortcut did not meaningfully affect our results, we conducted a simulation in a scenario with genetic nurture where we increased the number of SNPs from 5,000 to 100,000 and the GWAS discovery sample to 100,000. We hold the discovery sample fixed at *N* = 16, 000. In turn, we vary the level of assortative mating (*AM* = 0 or 1).

Figures 8 shows a design without assortative mating, and a design with assortative mating of 1, respectively. As can be seen, when holding constant the GWAS discovery sample size, and without assortative mating, Figure 8a shows that both ORIV as well as the PGI-RC produce a consistent estimator, whereas a meta-analysis PGI is substantially biased. In case of strong assortative mating, Figure 8a again shows that the PGI respository correction is more vulnerable than ORIV in overestimating the true coefficient when the outcome is subject to assortative mating. Figure 8b shows that ORIV only slightly underestimates the direct genetic effect within-families irrespective of the level of assortative mating. Overall, the same patterns arise as in our benchmark scenarios, suggesting that the shortcut to proportionally reducing the number of SNPs and the GWAS sample size does not meaningfully affect our conclusions.

**Figure 8.**
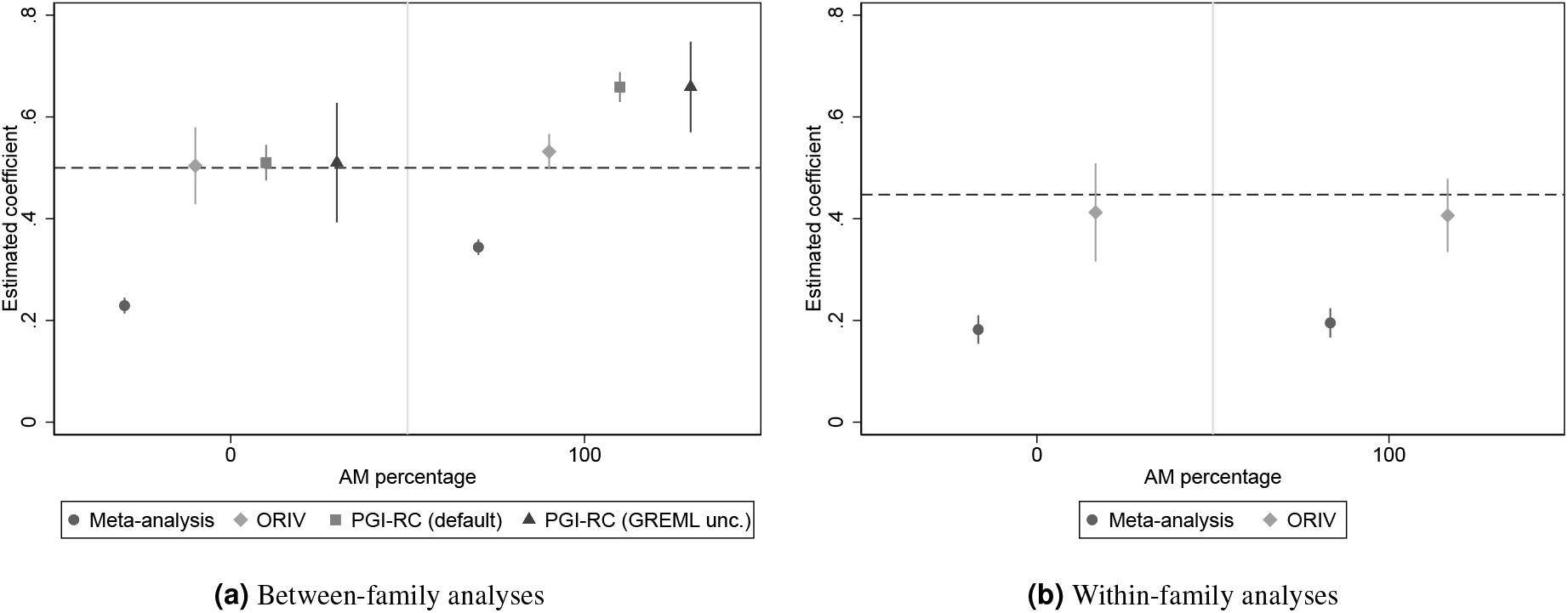
Estimated coefficients for the Polygenic Index (PGI) and their corresponding 95% confidence intervals using meta-analysis (circles), Obviously Related Instrumental Variables (ORIV, rhombuses), the default PGI-RC procedure (squares), and the PGI-RC procedure taking into account uncertainty in the GREML estimates (triangles) in a scenario with genetic nurture, varying assortative mating between 0 and 1, holding constant the prediction sample at *N* = 16, 000, and holding constant the GWAS discovery sample at *N* = 100, 000 and the number of SNPs also at *M* = 100, 000. Dashed line represents the true coefficient. The simulation results are based on 10 replications.

## D Data

The empirical analysis uses siblings in the UK Biobank (UKB) (29) as a prediction sample for constructing the polygenic indices (PGIs). The GWAS discovery sample excludes these full siblings and their relatives up to the 3rd degree to ensure that the discovery and the prediction samples are independent from each other. The analyses were restricted to individuals of European ancestry only. For this purpose, we excluded individuals with a value larger than 0 on the first principal component of the genetic data. We also filtered based on the self-reported ethnicity. Specifically, we only include individuals if their ethnic background is “White”, “British”, “Irish”, or “Any other white background”.

### D.1 Relatedness

The UKB provides a kinship matrix, which contains the genetically identified degree of relatedness for pairs of UKB participants related up to a third degree or closer. The kinship coefficient is constructed using KING software (72). We use the following values of the kinship coefficient corresponding to each degree of relatedness: monozygotic twins (*>* 0.354), first degree parent-child or full siblings (0.177-0.354), and second and third degree relatives and cousins (0.044-0.177). In order to separate parent-child pairs from siblings, we use the identity-by-state (*IBS*_0_) coefficient, which measures genetic similarity in terms of the fraction of markers for which the related individuals do not share alleles (29). Given their kinship coefficient, parent-child pairs have *IBS*_0_ *<* 0.0012 and full siblings have *IBS*_0_ *>* 0.0012. After classifying all relationships, we separated those who are related to the siblings up to the third degree other than siblings themselves, i.e., parents of siblings and cousins of siblings (*N* = 14, 947), and excluded those individuals from the GWAS discovery sample along with the full siblings. This ensures that our prediction sample of siblings is unrelated to the GWAS discovery sample. Importantly, in the GWAS discovery sample there are still some related (non-sibling) individuals. For this reason, we adjust for genetic relatedness in the GWASs. In particular, we kept one random cousin from the cousin clusters before randomly splitting the samples for GWAS.

### D.2 Phenotypes

We follow the literature (see e.g. (35), (12), (34)) and convert an individuals’ highest self-reported educational qualification to equivalent years of education using the International Standard Classification of Education (ISCED). The resulting *years of education* phenotype ranges from 7 to 20, where College or University degree is equivalent to 20 years, National Vocational Qualification (NVQ), Higher National Diploma (HND), or Higher National Certificate (HNC) to 19 years, other professional qualifications to 15 years, having an A or AS levels or similar to 13 years, O levels, and (General) Certificate of Secondary Education ((G)CSE) to 10 years. If “none of the above” is selected, then the lowest level of 7 years is assigned. For *height*, we use the standing height (in centimeters) of participants measured as a part of the anthropometric data collection at the UKB assessment center. For both phenotypes, we impute the missing values in the first wave with the available information from the two follow up measurements in the UKB.

### D.3 GWAS

We perform the GWAS on the UKB discovery sample for educational attainment (*N* = 389, 419) and for height (*N* = 391, 931), where both exclude siblings and their relatives (*N* = 56, 450). For the split-sample PGIs, we first removed all remaining parent-child pairs (*N* = 5, 084) and all cousins except one from each cousin cluster (*N* = 44, 326). Thereafter, we split the unrelated discovery sample randomly into two parts of 170, 004 for educational attainment and 170, 937 for height. We use the fastGWA approach (43) as implemented in Genome-wide Complex Trait Analysis (GCTA), which applies mixed linear modeling (MLM) to genetic data and requires the following steps. First, based on the relatedness matrix, we generate a sparse genetic relatedness matrix (GRM). We then use the sparse GRM, the SNP data, and the respective phenotype file for height and educational attainment to run the GWAS. Each phenotype file contains a measure of the phenotype residualized with respect to month and year of birth, gender, interaction of birth year and gender, genotyping batch, and the first 40 principal components (PCs) of the genetic relationship matrix. Additional quality control includes cleaning the data with respect to exclusion of individuals who withdrew consent, with bad genotyping quality, with putative sex chromosome aneuploidy, whose second chromosome karyotypes are different from XX or XY, with outliers in heterozygosity, with missing gender or self-reported gender mismatching with the genetically identified gender, individuals of non-European ancestry, or with missing information on any of the former criteria, on the phenotype or on any of the control variables. Further, we perform quality control on the resulting GWAS summary statistics using the EasyQC tool (73).

### D.4 Meta-Analysis

We meta-analyze our own GWAS summary statistics obtained with UKB data with the 23andMe summary statistics for the educational attainment PGI and with the GIANT 2014 summary statistics (41) for the height PGI using the software package METAL (70). While the GIANT summary statistics are publicly available, the 23andMe summary statistics are only available through 23andMe to qualified researchers under an agreement with 23andMe that protects the privacy of the 23andMe research participants. For more information, visit https://research.23andme.com/collaborate/\#dataset-access/. 23andMe research participants were included in the analysis on the basis of consent status as checked at the time data analyses were initiated. The research participants provided informed consent and participated in the research online, under a protocol approved by the external AAHRPP-accredited IRB, Ethical & Independent Review Services (E&I Review).

### D.5 Polygenic indices

We construct the PGIs by accounting for the linkage disequilibrium (LD), i.e., the non-random correlations between SNPs at various loci of a single chromosome, using the LDpred tool (42), version 1.06, and Python, version 3.6.6. LDpred is a Python based package that corrects the GWAS weights for LD using a Bayesian approach. We follow steps as summarized by (74): (i) coordinate the base and the target files, (ii) compute the LD adjusted weights, and (iii) construct the polygenic index with PLINK (75) using the LD weighted GWAS summary statistics. As LD reference panel, we use 30,000 randomly selected unrelated individuals of EU ancestry in the UKB who passed the quality control protocol. The PGIs are based on approximately 1 million HapMap3 SNPs assuming an infinitesimal model (i.e., we do not apply *p*-value thresholds). More details can be found in Table 11. The final prediction sample consists of *N* = 35, 282 siblings.

**Table 11.** Construction details Polygenic Indices.

| Trait | GWAS discovery sample | GWAS sample size | #SNPs in PGI |
| --- | --- | --- | --- |
| EA | UKB | 389,419 | 1,065,147 |
|  | UKB (Split sample 1) | 170,004 | 1,065,147 |
|  | UKB (Split sample 2) | 170,004 | 1,065,147 |
|  | 23andMe | 365,536 | 1,057,144 |
|  | UKB + 23andMe | 754,955 | 1,065,061 |
| Height | UKB | 391,931 | 1,065,146 |
|  | UKB (Split sample 1) | 170,937 | 1,065,145 |
|  | UKB (Split sample 2) | 170,937 | 1,065,145 |
|  | GIANT | 253,280 | 969,902 |
|  | UKB + GIANT | 654,211 | 1,065,063 |

## E Robustness empirical results

### E.1 Diastolic blood pressure

In Table 12, we present the empirical results for Diastolic Blood Pressure (DBP), where we used only a split-sample IV approach on basis of the UKB non-sibling subsample as described in Supplementary Information D. In line with what can be expected because of the weak assortative mating on this phenotype (76), we do not find strong differences across the between-family (top panel) and within-family (bottom panel) results. Similar as for the other outcomes we studied, a meta-analysis increases the standardized coefficient and increases the predictive power (here, from ∼ 4% to ∼ 6%). Applying ORIV further boosts the standardized effect by another 27%. The estimate obtained by the PGI-RC method is even higher, yet is surrounded by larger confidence intervals, especially when taking into the uncertainty of the GREML estimate. Within-families, ORIV again substantially boosts the estimated coefficient by 27%, clearly outperforming an OLS regression on basis of the meta-analysis PGI.

**Table 12.** Results of the OLS and IV regressions explaining (residualized and standardized) Diastolic Blood Pressure (DBP).

|  | OLS<br>(UKB 1) | OLS<br>(UKB 2) | OLS<br>(Meta-analysis) | ORIV<br>(Split-sample) | PGI-RC<br>(Default) | PGI-RC<br>(GREML unc.) |
| --- | --- | --- | --- | --- | --- | --- |
| **Between-family results** |  |  |  |  |  |  |
| Polygenic index | 0.209***<br>(0.006) | 0.204***<br>(0.006) | 0.249***<br>(0.006) | 0.318***<br>(0.006) | 0.366***<br>(0.008) | 0.366***<br>(0.029) |
| First-stage estimate |  |  |  | 0.422***<br>(0.005) |  |  |
| First-stage $F$-statistic | | | | 6541.31 | | |
| Incremental $R^2$ | 4.3% | 4.2% | 6.2% | | | |
| Family fixed effects | NO | NO | NO | NO | NO | NO |
| $N$ | 31,977 | 31,977 | 31,977 | 31,977 | 31,977 | 31,977 |
| **Within-family results** |  |  |  |  |  |  |
| Polygenic index | 0.189***<br>(0.010) | 0.204***<br>(0.010) | 0.243***<br>(0.010) | 0.308***<br>(0.014) |  |  |
| First-stage estimate |  |  |  | 0.415***<br>(0.006) |  |  |
| First-stage $F$-statistic | | | | 3936.66 | | |
| Incremental $R^2$ | 3.6% | 4.1% | 5.9% | | | |
| Family fixed effects | YES | YES | YES | YES |  |  |
| $N$ | 31,977 | 31,977 | 31,977 | 31,977 | | |

Notes: \* $p$ -value $< 0.10$ ; \*\* $p$ -value $< 0.05$ ; \*\*\* $p$ -value $< 0.01$ . In all regressions the dependent variable is residualized diastolic blood pressure (standardized to have mean 0 and standard deviation 1), where the residuals are obtained after a regression of diastolic blood pressure controlling for sex, year of birth, month of birth, sex interacted with year of birth, and the first 40 principal components of the genetic relationship matrix. Standard errors are robust and clustered at the family level, and in case of ORIV also at the individual level. OLS (UKB 1/2) refers to models with the PGI constructed using the split-sample UKB non-sibling (i.e., excluding all siblings and their relatives) sample. OLS (Meta-analysis) uses a PGI constructed using a meta-analysis of the two split-sample UKB non-sibling samples. ORIV (Split-sample) refers to a 2SLS estimation where the summary statistics derive from a random split of the UKB sample. PGI-RC refers to the PGI-RC method, where (Default) refers to the conventional application of the method, whereas (GREML unc.) refers to the case where we incorporate the uncertainty in the estimation of the SNP-based heritability.

### E.2 The between-family and within-family component of the PGI

The correlation between the independent PGIs reflects the measurement error in the PGI of the between-family component. There will be some bias in the within-family estimates if the measurement error of the within-family signal is considerably different from the measurement error of the between-family signal. We therefore computed the correlation between deviations of the PGI from the family means in the UKB, and use this as the scaling factor in the within-family analysis.^11^ The results were very similar compared to the main results. For EA, the within-family two-sample and split-sample coefficients were estimated as 0.179 (0.012) and 0.161 (0.012), respectively. For comparison, the coefficients are 0.184 (0.012) and 0.170 (0.013) in Table 2. For height, the within-family two-sample and split-sample ORIV coefficients were estimated as 0.604 (0.008) and 0.556 (0.009), respectively. In Table 3, the coefficients are 0.614 (0.008) and 0.571 (0.009). Full results for this robustness check are available upon request from the authors. These findings suggest that the measurement error across between- and within-family signals is comparable and does not represent a relevant threat to the application of ORIV for these traits.

### E.3 Comparing ORIV with regular IV

In this section we compare the ORIV estimator to the regular IV estimator suggested by DiPrete et al. (26) for our between-family empirical analyses regarding EA (Table 13) and height (Table 14). To obtain the regular IV estimator, a researcher needs to select which PGI to use as instrumental variable. The Obviously-Related Instrumental Variables (ORIV) approach (28) overcomes the arbitrary choice of selecting one PGI as the independent variable and the other PGI as IV.

**Table 13.** Results of the IV regressions explaining (residualized and standardized) educational attainment.

|  | IV<br>(UKB) | IV<br>(23andMe) | ORIV<br>(2-sample) | IV<br>(UKB 1) | IV<br>(UKB 2) | ORIV<br>(Split-sample) |
| --- | --- | --- | --- | --- | --- | --- |
| Polygenic index | 0.309***<br>(0.008) | 0.366***<br>(0.008) | 0.337***<br>(0.007) | 0.322***<br>(0.008) | 0.325***<br>(0.008) | 0.323***<br>(0.007) |
| *N* | 35,282 | 35,282 | 35,282 | 35,282 | 35,282 | 35,282 |

Notes: \* $p$ -value < 0.10; \*\* $p$ -value < 0.05; \*\*\* $p$ -value < 0.01. In all regressions the dependent variable is residualized EA (standardized to have mean 0 and standard deviation 1), where the residuals are obtained after a regression of height on controlling for sex, year of birth, month of birth, sex interacted with year of birth, and the first 40 principal components of the genetic relationship matrix. Family fixed effects are not included. Standard errors are robust and clustered at the family level, and in case of ORIV also at the individual level. IV (UKB) refers to a 2SLS estimation using PGI on the basis of the UKB instrumented by the PGI constructed using the 23andMe summary statistics, while IV (23andMe) is a 2SLS regression where the roles are reversed. ORIV (2-sample) refers to a 2SLS estimation using the PGIs from the UKB non-sibling sample and Giant as instrumental variables for each other. IV (UKB1(2)) refers to a 2SLS estimation using the split-sample UKB PGI 1(2) is used as IV for UKB PGI 2(1). ORIV (Split-sample) refers to a 2SLS estimation where the summary statistics derive from a random split of the UKB sample.

**Table 14.** Results of the IV regressions explaining (residualized and standardized) height.

|  | IV<br>(UKB) | IV<br>(GIANT) | ORIV<br>(2-sample) | IV<br>(UKB 1) | IV<br>(UKB 2) | ORIV<br>(Split-sample) |
| --- | --- | --- | --- | --- | --- | --- |
| Polygenic index | 0.571***<br>(0.006) | 0.734***<br>(0.008) | 0.652***<br>(0.006) | 0.625***<br>(0.007) | 0.634***<br>(0.007) | 0.617***<br>(0.007) |
| *N* | 35,282 | 35,282 | 35,282 | 35,282 | 35,282 | 35,282 |

*Notes:* \* $p$ -value < 0.10; \*\* $p$ -value < 0.05; \*\*\* $p$ -value < 0.01. In all regressions the dependent variable is residualized height (standardized to have mean 0 and standard deviation 1), where the residuals are obtained after a regression of height on controlling for sex, year of birth, month of birth, sex interacted with year of birth, and the first 40 principal components of the genetic relationship matrix. Family fixed effects are not included. Standard errors are robust and clustered at the family level, and in case of ORIV also at the individual level. IV (UKB) refers to a 2SLS estimation using PGI on the basis of the UKB instrumented by the PGI constructed using the GIANT summary statistics, while IV (GIANT) is a 2SLS regression where the roles are reversed. ORIV (2-sample) refers to a 2SLS estimation using the PGIs from the UKB non-sibling sample and GIANT as instrumental variables for each other. IV (UKB1(2)) refers to a 2SLS estimation using the split-sample UKB PGI 1(2) is used as IV for UKB PGI 2(1). ORIV (Split-sample) refers to a 2SLS estimation where the summary statistics derive from a random split of the UKB sample.

The results show that when using summary statistics from two different sources (UKB and 23andMe in the case of EA; UKB and GIANT in the case of height) the IV estimates can vary quite strongly and can be significantly different from each other. In these cases, rather than making an arbitrary choice for one or the other, it seems more appropriate to apply ORIV, where all information is used. In case of the split-sample PGIs, the ORIV and IV point estimates do not diverge as much. Still, even here, ORIV would be recommended as the precision of these estimates is slightly better.

## Footnotes

1 Young et al. (31) developed Related Disequilibrium Regression (RDR) to estimate heritability that avoids confounding from indirect genetic effects. The method however requires genetic data from unrelated individuals and both of their parents, samples of which are currently very rare. Moreover, similar to other heritability estimators, it assumes the absence of gene–environment interplay while empirical evidence shows this may be important (e.g., 19; 20).

2 To verify that our results are not sensitive to the shortcut of downsizing both *M* and *N*_*GWAS*_ for computational reasons, we also analyze an additional setting with *M* = 100, 000 and *N*_*GWAS*_ = 100, 000 (10 runs). The results remain very similar, see Supplementary Information C.2.

3 It is also lower than the reported *R*^2^ of around 12% in (35), but that study used a larger GWAS meta-analysis sample to construct the PGI and different prediction samples.

4 Since the total genetic effect is the sum of the direct and indirect genetic effects, our within-family estimates also implicitly estimate an upper bound on the indirect genetic effect. Comparing the between- and within-family estimates, our analysis suggests that this upper bound is approximately (0.337 − 0.184)*/*0.337 = 45% for EA and approximately (0.652 − 0.614)*/*0.652 = 6% for height. As such, we can compare this to earlier studies that estimated indirect genetic effects. For example, while one study (47) estimates that the indirect genetic effect for EA is approximately 30% of the total genetic effect, a second study (18) estimates that up to 60% of the total genetic effect could be indirect. Our estimate of 45% is exactly in the middle of these estimates, and thus represents a tightened upper bound on the indirect genetic effect on EA.

5 They denote the factor by *ρ*, but to avoid confusion with our definition of *ρ* being the correlation between the two PGIs, we do not use this symbol here.

6 In rare cases, in a very small prediction sample, the estimated SNP-heritability in our simulations is estimated to be very small. This results in a huge standard error as can be seen in (14). To avoid that these rare cases have a large influence on our mean comparisons, we ignore runs that produce a SNP-based heritability lower than 0.01.

7 Since environments differ, the best linear genetic predictor (i.e., the true latent PGI) may differ across samples. This would imply that the genetic correlation between the two samples would be lower than 1 for a particular outcome variable. This can be tested, for example with LDSC (66). LDSC can also be used to estimate the cross-trait LDSC intercept. With a trend towards ever-larger GWAS meta-analyses and reuse of genetic data, it is important to verify that GWAS summary statistics used to create the two independent PGIs are not computed from partially overlapping samples (or from sets of individuals that are related to each other). Using two PGIs based on such related GWAS summary statistics would be in violation of the ORIV assumptions. The cross-trait LDSC intercept can be used to test for sample overlap, although the precise threshold depends on the GWAS samples sizes and the heritability of the trait (67). In our study, the LDSC cross-trait intercepts do not cross the critical threshold (results available upon request). Moreover, in the UKB split-sample analyses, the GWAS samples do not contain related individuals (see Supplementary Information D).

8 In rare cases for a very small GWAS discovery sample, a simulation run would produce a negative correlation between the two PGIs. We ignore these runs in the resulting averages.

9 In Supplementary Information E.2 we show that our ORIV estimates are not biased when applying the between-family correlation between PGIs in the within-family analyses.

10 Technically, for Figure 1a we would require a design without genetic nurture, but since the SNP-based heritability is held constant and the Figure refers to a between-family setting, in practice this does not matter.

11 We thank Elliot Tucker-Drob for this suggestion.

